# Identification of mutations that cooperate with defects in B cell transcription factors to initiate leukemia

**DOI:** 10.1101/2020.11.25.398966

**Authors:** Lynn M. Heltemes-Harris, Gregory K. Hubbard, Rebecca S. La Rue, Sarah A. Munro, Todd P. Knudson, Rendong Yang, Christine M. Henzler, Timothy K. Starr, Aaron L. Sarver, Steven M. Kornblau, Michael A. Farrar

## Abstract

The transcription factors EBF1 and PAX5 are frequently mutated in B cell acute lymphoblastic leukemia (B-ALL). We demonstrate that *Pax5*^+/-^ *x Ebf1*^+/-^ compound heterozygous mice develop highly penetrant leukemia. Similar results were seen in *Pax5*^+/-^ *x Ikzf1*^+/-^ and *Ebf1*^+/-^ *x Ikzf1*^+/-^ mice for B-ALL, or in *Tcf7*^+/-^ *x Ikzf1*^+/-^ mice for T cell leukemia. To identify genetic defects that cooperate with *Pax5* and *Ebf1* compound heterozygosity to initiate leukemia, we performed a Sleeping Beauty (SB) transposon screen that identified cooperating partners including gain-of-function mutations in *Stat5 (∼65%)* and *Jak1(∼68%)*, or loss-of-function mutations in *Cblb* (61%) and *Myb (32%)*. These findings underscore the role of JAK/STAT5 signaling in B cell transformation and demonstrate unexpected roles for loss-of-function mutations in *Cblb* and *Myb* in leukemic transformation. RNA-Seq studies demonstrated upregulation of a PDK1>SGK3>MYC pathway; treatment of *Pax5*^+/-^ *x Ebf1*^+/-^ leukemia cells with PDK1 inhibitors blocked proliferation in vitro. Finally, we identified conserved transcriptional variation in a subset of genes between human leukemias and our mouse B-ALL models. Thus, compound haploinsufficiency for B cell transcription factors likely plays a critical role in transformation of human B cells and suggest that PDK1 inhibitors may be effective for treating patients with such defects.

## Introduction

Heterozygous deletions or loss of function mutations in a number of B cell transcription factors are a common feature of human B cell acute lymphoblastic leukemia (ALL)[1]. This is clearly evident for three transcription factors - EBF1, PAX5 and IKZF1[1, 2]. Interestingly, alterations involving these transcription factors commonly occur together[1, 3]. This is particularly pronounced in BCR-ABL^+^ leukemia in which 50% of leukemias with *IKZF1* deletions also have mutations affecting *Pax5* expression or function[4]. Therefore, an important question is whether compound haploinsufficiency for these transcription factors drives transformation and which combinations of transcription factors can promote transformation. Finally, the genetic alterations that cooperate with haploinsufficiency for these transcription factors to drive transformation have also not been comprehensively elucidated.

To address the questions above, we generated a set of mice that exhibited compound haploinsufficiency for various combinations of *Ebf1, Pax5, Ikzf1, Cebpa*, and *Tcf7*. Herein, we demonstrate that *Pax5*^+/-^ *x Ebf1*^+/-^, *Pax5*^+/-^ *x Ikzf1*^+/-^, and *Ebf1*^+/-^ *x Ikzf1*^+/-^ mice all generated B cell leukemia, while *Tcf7*^+/-^ *x Ikzf1*^+/-^ mice generated T cell leukemia. Furthermore, we used a SB Transposon screen to identify mutations that cooperate with *Pax5*^+/-^ *x Ebf1*^+/-^ compound haploinsufficiency to promote transformation. Our findings document the key role that compound haploinsufficiency for critical transcription factors plays in leukemia transformation and identify mutations that cooperate with such alterations to initiate transformation.

## MATERIALS AND METHODS

### Mice and Cells

All mice have been previously described [5-10]; the University of Minnesota IACUC approved all animal experiments. Mice were monitored for up to 400 days for leukemia. Spleen, lymph nodes, and bone marrow were isolated from tumor-bearing mice and used for further experiments.

### RNA-Seq Analysis

RNA-seq was performed on total RNA extracted from either column purified progenitor B control cells (C57Bl/6, *Pax5*^+/-^*x Ebf1*^+/-^, *Pax5*^+/-^ *or Ebf1*^+/-^) or leukemic cells from lymph nodes of tumor-bearing mice using a RNeasy Plus kit (Qiagen). Fifty-three barcoded TruSeq RNA v2 libraries were created and sequenced on a HiSeq 2500. A second set of data were used for variant calling analysis. Eight barcoded libraries were sequenced on the HiSeq 2000 to produce 100 bp paired end reads.

### Systematic identification of gene clusters

Phase_2 BCCA and St Jude (2016) and phase_3 St Jude ALL(2018) Human mRNA RNA-Seq data were downloaded from https://ocg.cancer.gov/programs/target/data-matrix on April 23 2019 (dbGaP accession phs000218.v22.p8). The Value of 0.1 was added to each value the data was mean centered and log transformed. A SD cutoff was used to identify ∼8500 genes in each of three datasets. Unsupervised hierarchical clustering was used to define sets of genes which were defined by average linkage correlation > 0.2 and more than 150 members. Statistical enriched Gene cluster memberships across clusters were defined by Fisher exact test to identify “common” clusters across datasets. For the mouse data, the tumors were treated in a similar fashion except an SD cutoff of 1.0 was used. Statistical enriched gene cluster memberships across clusters were defined by Fisher exact test to identify “common” clusters across datasets using gene name matching.

### Data Analysis

Data was analyzed using Prism 8 (Graphpad). A Shapiro-Wilk test was used to determine normality of all data. Unpaired data that passed normality was analyzed by an ordinary one-way ANOVA with Holm-Sidak’s multiple comparison test or by an unpaired t-test; data that failed normality were analyzed using an unpaired Kruskal-Wallace test with Dunn’s multiple comparison test. Kaplan-Meier Survival curves were analyzed by Log-rank (Mantel-Cox) Test. Integrated Genomics Viewer was used to view aligned sequences (Broad Institute).

### Accession Numbers

RNA-Seq data was deposited with GEO and is accessible through GEO Series accession number GSE148680 (https://www.ncbi.nlm.nih.gov/geo/query/acc.cgi?acc=GSE148680).

### Supplemental methods

Supplemental Methods section includes detailed protocols of cell lines and culture conditions, NGS, flow cytometry, qPCR, western blotting, Gene Set Enrichment Analysis, SB Mutagenesis, Transposon Insertion Analysis, Reverse Phase Proteomics and Inhibitor Assay.

## RESULTS

### Reduced expression of transcription factors critical for lymphocyte development leads to leukemia

To explore whether compound haploinsufficiency for *Ebf1* and *Pax5* leads to B cell transformation, we generated *Pax5*^+/-^ *x Ebf1*^+/-^ mice. As shown in Figure 1A, ∼67% of *Pax5*^+/-^ *x Ebf1*^+/-^ mice develop leukemia with a mean survival of 296 days. Flow cytometry analysis from bone marrow, lymph nodes and spleens of these mice demonstrated that leukemias resemble progenitor-B cells with a B220^+^CD19^+^IgM^-^ phenotype (Fig. 1B) and also express pre-BCR, CD43, IL7R, TSLPR, c-KIT, AA4.1 and CD25 confirming their progenitor-B cell like phenotype (Sup Fig. 1). Although both male and female mice developed leukemia in this model, female mice exhibited greater penetrance (97% versus 71% at 400 days) and reduced median survival (265 days vs 298 days, p= 0.005; Sup Fig. 2). The effect of reduced expression of *Pax5* and *Ebf1* on transformation was not limited to this combination of transcription factors as similar results were observed in *Pax5*^+/-^ *x Ikzf1*^+/-^ and *Ebf1*^+/-^ *x Ikzf1*^+/-^ mice (Fig. 1A). Compound haploinsufficiency for all three transcription factors (*Pax5*^+/-^ *x Ebf1*^+/-^ *x Ikzf1*^+/-^ mice) resulted in 100% penetrance of leukemia and much shorter mean survival (202 days). We previously reported that *Pax5*^+/-^ or *Ebf1*^+/-^ mice do not develop leukemia[11]. In contrast, *Ikzf1*^+/-^ mice do develop leukemia with low penetrance (Fig. 1A)[12, 13]; however, these were always T cell leukemias (Fig. 1C). Deleting one copy of *Pax5* and *Ebf1* not only increased the frequency of B cell leukemias in *Ikzf1*^+/-^ mice (none to ∼40%), but surprisingly, also resulted in a dramatic increase in T cell leukemias (∼5% in *Ikzf1*^+/-^ mice versus ∼35% in *Pax5*^+/-^ *x Ebf1*^+/-^ *x Ikzf1*^+/-^ mice)(Fig. 1C,D). Thus, although PAX5 and EBF1 are only expressed in B cells, reduced expression of these two transcription factors paradoxically also promoted T cell leukemia.

**Figure 1.**
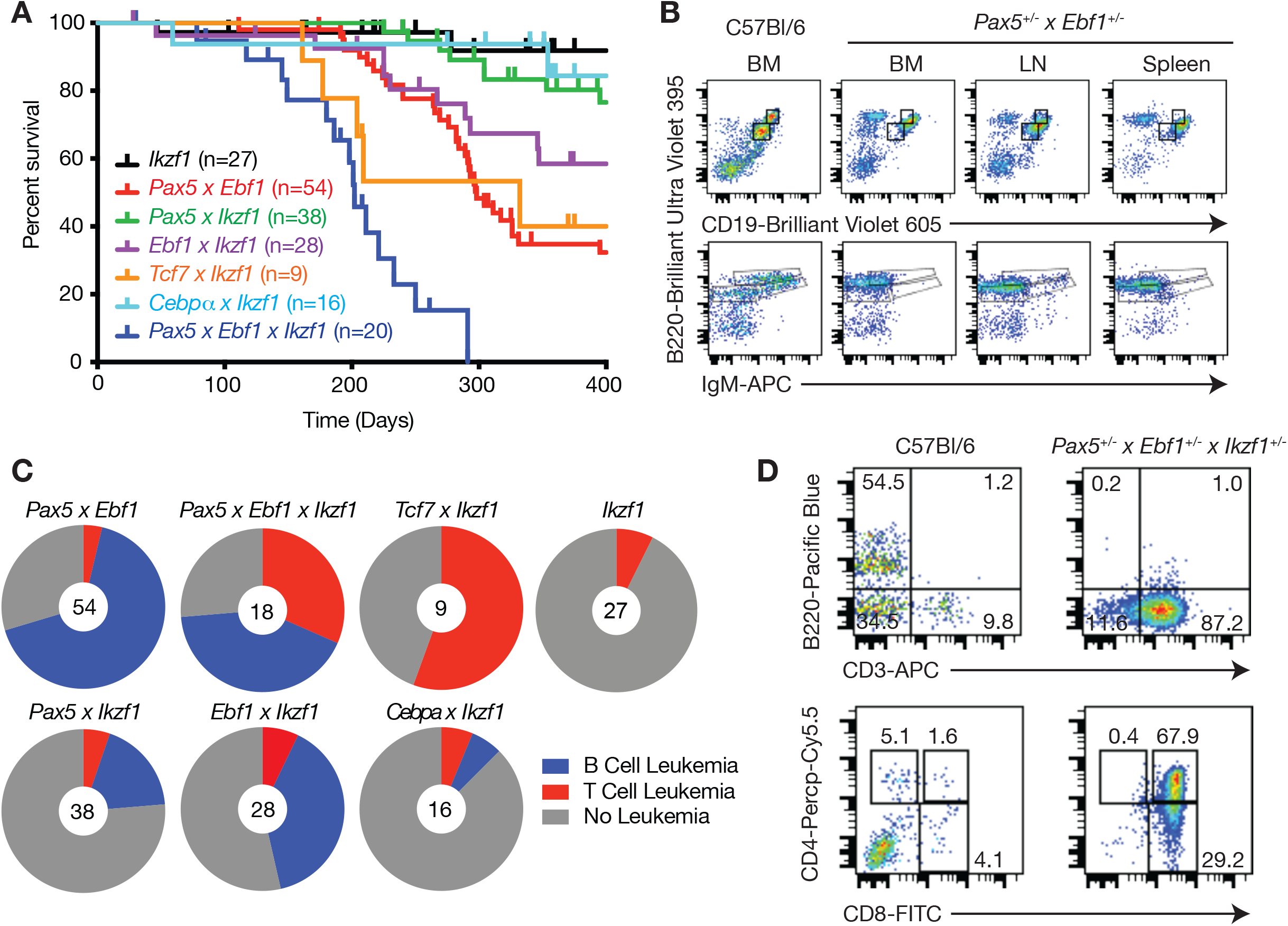
Compound haploinsufficiency for transcription factor genes drives B cell or T cell leukemia. **A** Kaplan-Meier survival analysis of mice of the indicated genotype. **B** Flow cytometric analysis of control C57Bl/6 bone marrow (BM) cells or bone marrow, lymph node (LN), and spleen cells from *Pax5*^+/-^ *x Ebf1*^+/-^ leukemic mice. Representative flow cytometric analysis of B220, CD19, and IgM expression is shown. Doublets were gated out and a lymphocyte gate was set based on side and forward scatter properties. All gates shown are based on bone marrow isolated from control C57Bl/6 mice. **C** Pie charts showing the number of leukemias from each genotype that were either of the B cell (Blue) or T cell (Red) phenotype; grey represents mice that either failed to develop leukemia or developed mixed lineage leukemia (grey). **D** Flow cytometric analysis of bone marrow cells from control C57Bl/6 and *Pax5*^+/-^ *x Ebf1*^+/-^ *x Ikzf1*^+/-^ leukemic mice. Representative flow cytometric analysis of B220, CD3, CD4, and CD8 expression on bone marrow cells is shown. Doublets were gated out and a lymphocyte gate was set based on side and forward scatter properties. All gates shown are based on bone marrow isolated from control C57Bl/6 mice.

**Figure 2.**
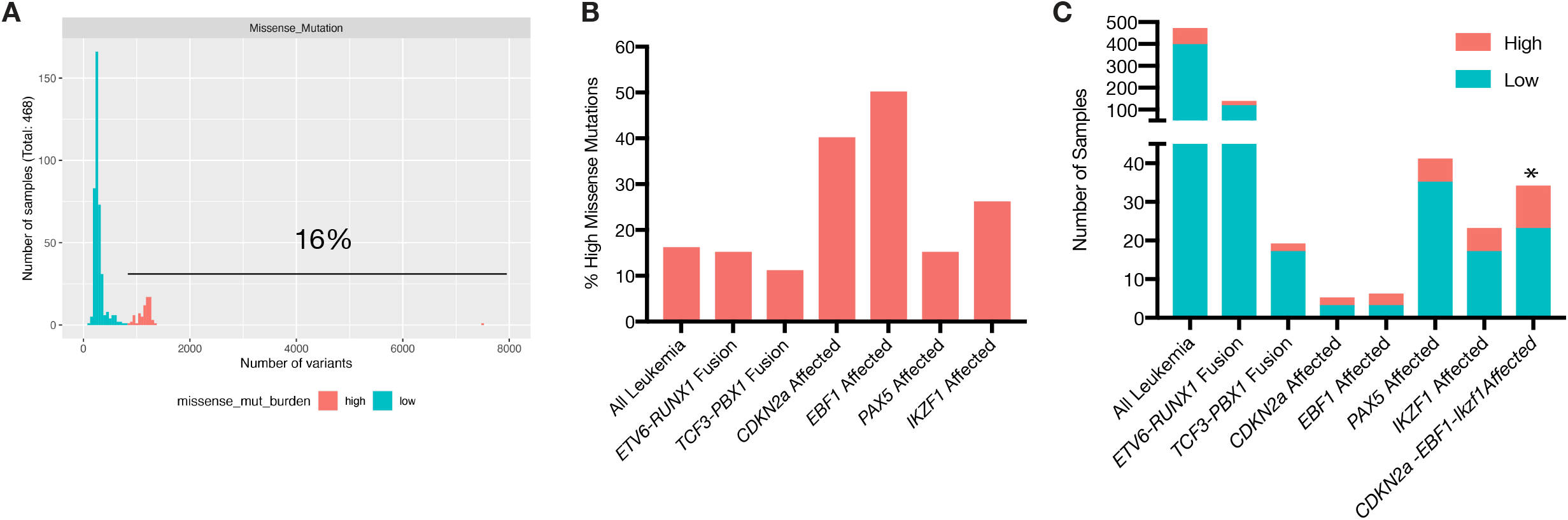
Missense mutations in primary human ALL. **A** Histogram showing the distribution of missense mutations per sample across 486 primary human B-ALL samples from the TARGET phase-2 study. SNV/indel variants were called across all genes by MuTect2. Samples were classified as high or low mutation burden based on the natural break in the bimodal distribution (low < 669, high > 863 variants/sample). The percentage of samples in the high category is labeled inside the plot. **B** Bar graph summarizing the percentage of samples with high mutation burden compared to low for all B-ALL samples or various subsets. **C** Bar graph summarizing the total number of primary human B-ALL samples displaying high or low missense mutation burden. Samples affected by loss of function alleles in various genes (*CDKN2a, EBF1, PAX5, and IKZF1*) or contain certain gene-fusions (*ETV6-RUNX1, TCF3-PBX1*) were shown as subsets. * p<0.0114 Fisher’s exact test.

We next examined whether compound haploinsufficiency for lineage determining transcription factors was a general mechanism that could promote transformation of multiple cell lineages. To this end, we generated *Tcf7*^+/-^ *x Ikzf1*^+/-^ mice, as *Tcf7*, which encodes TCF1, and *Ikzf1*, are both required for T cell development[8, 9]. In addition, we generated *Cebpa*^+/-^ *x Ikzf1*^+/-^ mice, as *Cebpa* and *Ikzf1* are both involved in myeloid cell development[14]. *Cebpa*^+/-^ *x Ikzf1*^+/-^ mice did not develop myeloid leukemia and the rate of T cell leukemia in *Cebpa*^+/-^ *x Ikzf1*^+/-^ mice was no higher than that observed for *Ikzf1*^+/-^ mice. Thus, not all combinations of transcription factor haploinsufficiency promote transformation. However, *Tcf7*^+/-^ *x Ikzf1*^+/-^ mice developed T cell leukemias with high penetrance, comparable to that seen for B cell leukemias in *Pax5*^+/-^ *x Ebf1*^+/-^ mice (Fig. 1A,C). Thus, compound haploinsufficiency for lineage defining transcription factors can promote transformation in multiple cell lineages and may underlie many types of leukemias.

### Genetic mutations that cooperate with Pax5 and Ebf1 heterozygosity to induce leukemia

Previous reports by Prasad and colleagues suggested that *Ebf1*^+/-^ mice have defects in DNA repair with decreased expression of *Rad51* and increased γH2AX foci[15]. These studies further claimed that defects in DNA repair resulted in increased mutation rates in *Pax5*^+/-^ *x Ebf1*^+/-^ leukemias and that this accounts, in part, for progenitor B cell transformation in those mice[15]. This suggestion is difficult to reconcile with the relatively low frequency of somatic mutations reported in human in B-ALL[16]. We re-examined this issue using *Ebf1*^+/-^ mice in our colony. In contrast to the previous study, we found no difference in *Rad51, Rad51AP or* γH2AX expression when examining log2 transformed FPKM values generated from two separate RNA-seq experiments (Sup. Fig. 3A). In fact, the low level of variation paralleled that observed for a panel of housekeeping genes (*B2m, Hprt*, and *Actb*; Sup. Fig. 3A). Since the previous studies compared progenitor-B cells from WT and *Ebf1*^+/-^ mice that had been cultured extensively in vitro we examined γH2AX expression in long-term cultured progenitor-B cells from WT and *Ebf1*^+/-^ mice; no significant difference was observed (Sup. Fig. 3B). Further, we examined γH2AX expression by flow cytometry in progenitor-B cells directly from the bone marrow of WT and *Ebf1*^+/-^ mice. Again, we found no significant difference in expression (Sup. Fig. 3C). Next, we examined whether genes involved in DNA repair were enriched in *Ebf1*^+/-^ cells by Gene Set Enrichment Analysis (GSEA) using our RNA-seq data. We saw a significant enrichment for DNA repair genes (Sup. Fig. 3D), although it is unclear whether this reflects a direct effect of EBF1 on genes involved in DNA repair or just a relative increase in cells stuck at a stage undergoing VDJ recombination, as there is significant overlap between genes involved in DNA damage response and VDJ recombination. Finally, we examined whether subsets of human B cell leukemias exhibited increased mutation rates and if so, whether they were enriched in those containing *Ebf1* mutations. As shown in figure 2A, ∼16% of B-ALLs obtained from the NIH TARGET ALL database showed significant levels of missense mutations. We broke the total B-ALL samples down into smaller subsets, characterized by *ETV6-RUNX1* or *TCF3-PBX1* translocations, or those with missense mutations in *CDKN2A, PAX5, EBF1* or *IKZF1*. Leukemias expressing the *ETV6-RUNX1* or *TCF3-PBX1* translocations, or *PAX5* missense mutations, were not enriched in hypermutated leukemias (Fig. 2B,C). In contrast, leukemias with missense mutations in *CDKN2A, EBF1* or *IKZF1* showed an increased percentage of leukemias with high numbers of missense mutations (Fig.2B). The number of *CDKN2A, EBF1*, or *IKZF1* samples was too small to assess whether the increased percentage of hypermutated leukemias was statistically significant. However, mutations in *EBF1, IKZF1* and *CDKN2A* are all enriched in *BCR-ABL*-like leukemias and when we pooled samples with these three mutations together there was a clear enrichment in samples with high numbers of missense mutations (Fig. 2C). In conclusion, leukemias with *EBF1* mutations may be preferentially found in hypermutated B-ALL, but this is not a feature restricted to *EBF1* as mutations in *CDKN2A* and *IKZF1* are also associated with this hypermutated phenotype.

**Figure 3.**
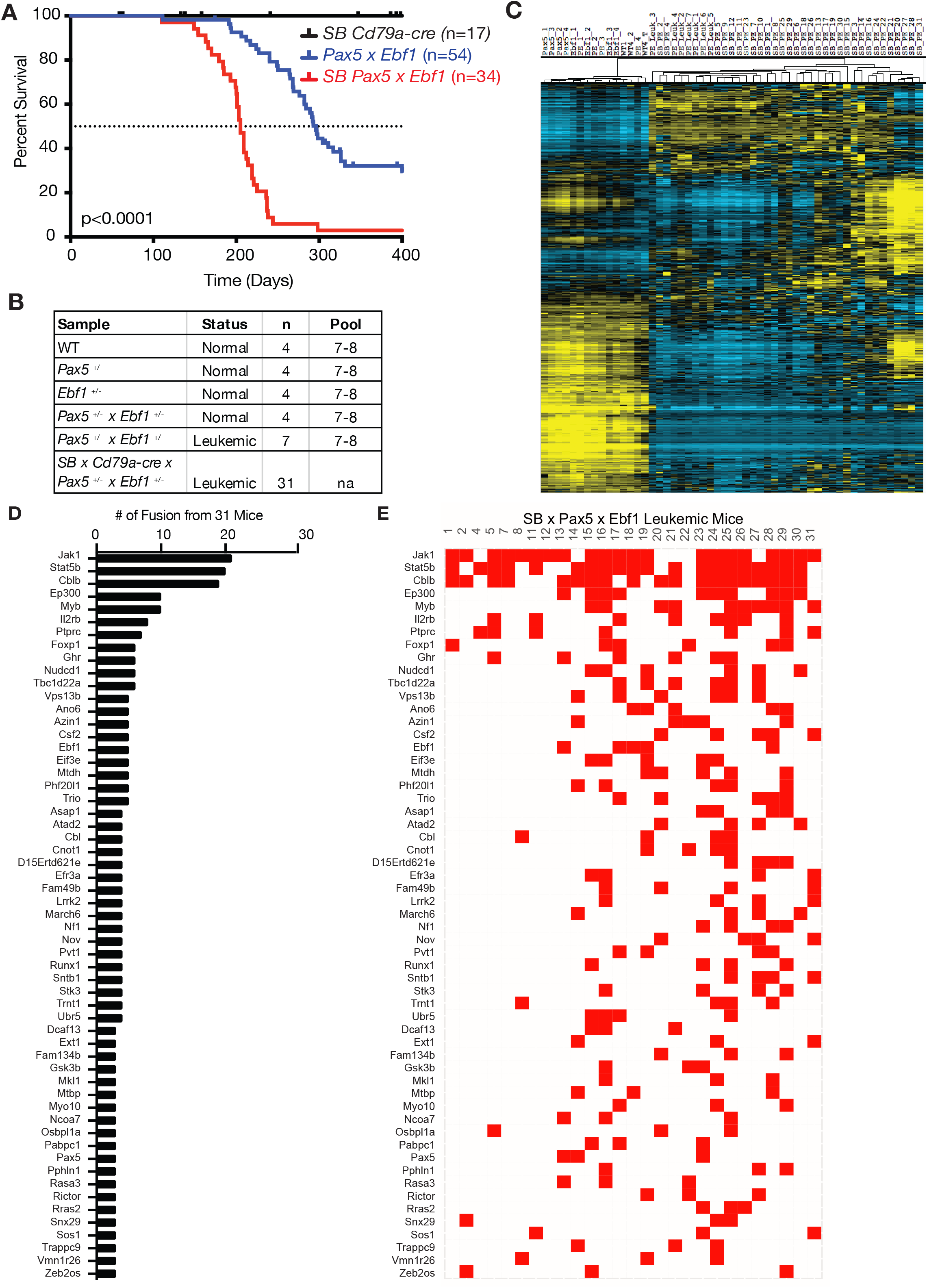
Sleeping Beauty mutagenesis screen to identify genes that cooperate with *Pax5* and *Ebf1* heterozygosity to induce leukemia. **A** Kaplan-Meier survival analysis of mice comparing *Pax5*^+/-^ *x Ebf1*^+/-^ leukemic mice (n=51) and SB *Pax5*^+/-^ *x Ebf1*^+/-^ (n=34) leukemic mice to control mice *SB x Cd79a-Cre* (n=17). P-value compares *Pax5*^+/-^ *x Ebf1*^+/-^ versus SB *Pax5*^+/-^ *x Ebf1*^+/-^ mice. **B** Table indicating all of the samples used in RNA-seq analysis. The table indicated the status, type and number of samples utilized for the RNA-Seq. Control samples represent progenitor B cell pools from 7-8 mice. **C** Hierarchical clustering of all leukemic and control samples. **D** Fusions identified from RNA-seq analysis of our sleeping beauty mutagenesis screen. This chart identifies recurrent insertions found in 27 of the 31 samples tested and indicate how many mice had each specific gene insertion. **E** Matrix analysis of individual mice by gene identified in fusion analysis.

To discover novel genes that cooperate with *Pax5* and *Ebf1* heterozygosity to induce B-ALL we employed a Sleeping Beauty (SB) transposon mutagenesis screen[17]. *Pax5*^+/-^ *x Ebf1*^+/-^ *x Cd79a-cre* mice were crossed to mice expressing the mutagenic transposon SB in a *Cd79a-Cre*-dependent, and hence B cell-specific, manner. We generated 34 mice that were heterozygous for both *Ebf1* and *Pax5* and expressed the mutagenic transposon. Mice were housed for up to 400 days to allow them to develop leukemia. We also included single heterozygous combinations (*Pax5*^+/-^*x Cd79a-Cre x SB and Ebf1*^+/-^ *x Cd79a-Cre x SB*) but neither of these cohorts developed leukemia within 400 days (data not shown). As seen in figure 3A, all of *Pax5*^+/-^*x Ebf1*^+/-^ *x Cd79a-Cre x SB* mice developed leukemia. Thus, the presence of the sleeping beauty transposon increased penetrance of leukemia from 67% to 100% and decreased the median age of death from 296 to 205 days. Thus, other genes mobilized or silenced by SB transposition clearly cooperate with *Pax5* and *Ebf1* heterozygosity to initiate transformation.

To identify which genes were targeted by the transposon, we performed RNA-seq analysis on 31 SB induced leukemias. The SB transposon contains a unique 5’ leader sequence with a splice donor site that allows for splicing into transcripts. In addition, the SB transposon also has a splice acceptor and SV40 polyA tail that allows for splicing of upstream exons to the SV40 poly A sequence, thereby allowing for premature termination. These unique 5’ SB sequences and 3’ SV40 polyA signal sequences can be identified by RNA-Seq as novel fusion proteins. This allowed us to map SB fusions and determine how transposon insertions altered specific gene expression[18]. We carried out RNA-Seq analysis on progenitor B cells (CD19^+^B220^+^Igκ/λ^-^) from WT, *Ebf1*^+/-^, *Pax5*^+/-^, and *Pax5*^+/-^ *x Ebf1*^+/-^ pre-leukemic mice (∼6-12 weeks of age), as well as spontaneous *Pax5*^+/-^ *x Ebf1*^+/-^ leukemias (Fig. 3B). Differential gene expression analysis showed that WT and pre-leukemic samples all clustered distinctly from the SB-induced and spontaneous leukemias. The spontaneous *Pax5*^+/-^ *x Ebf1*^+/-^ leukemias were interspersed among the SB-induced leukemias demonstrating that the SB-induced leukemias shared gene expression signatures with the spontaneous leukemias. Finally, the leukemias were clearly heterogenous with a number of distinct subsets harboring distinct gene signatures (Fig. 3C).

### RNA fusion analysis defines genes that cooperate with *Pax5 x Ebf1* heterozygosity to induce leukemia

To identify the targets of transposon mutagenesis, we performed an RNA-Seq-based analysis of transposon fusions to identify genes targeted in our screen. The fusion transcripts are detected either directly as unique gene fusions or can be imputed from paired end reads that have one end derived from SB and the second end derived from the target gene sequence (called bridging fusions)[18]. There were 758 unique gene fusions or bridging fusions that were used to identify recurrent fusion events in 27 of 31 leukemias. Figure 3D lists all the reoccurring RNA fusions identified in our screen. Consistent with the heterogeneity of the gene expression profiles in distinct B-ALL subsets (Fig 3C), many of the targeted genes were only found in a fraction of the leukemias (Fig. 3E). The most notable exception was that almost all leukemias had SB insertions involving either *Jak1* or *Stat5b* (Fig. 3E). SB RNA fusion analysis demonstrated that the SB 5’ leader UTR sequence typically fused to the first 1-4 coding exons of *Stat5b* or *Jak1* genes (Fig. 4A, Sup. 4A). This suggested that a full-length or nearly full-length coding transcript would be generated for both *Jak1* and *Stat5b. Stat5b* mRNA was expressed at significantly higher levels in leukemic samples harboring a SB transposon insertion (Fig. 4B,C). Consistent with data for *Stat5b* mRNA, there was a clear increase in the expression of STAT5 protein in samples with an SB insertion in the *Stat5b* gene (Fig. 4D-E). In contrast, the spontaneous *Pax5*^+/-^ *x Ebf1*^+/-^ leukemias did not exhibit significant increases in *Stat5b* expression (Fig. 4B,C). However, when we examined levels of phosphorylated STAT5 we found that *Pax5*^+/-^ *x Ebf1*^+/-^ leukemic cells expressed significantly higher levels of activated STAT5 than WT control mice, either directly ex vivo (Fig. 4F), or following in vitro stimulation with IL7 (Fig. 4G). This change represents a significant increase in phospho-STAT5 (p-STAT5) as it was equal to or higher than seen in mice expressing a constitutively active form of STAT5b in progenitor B cells (Fig. 4F,G)[19]. This result illustrated that there were increased levels of pStat5 expression in our leukemic cells. We next looked at known targets of STAT5b - *Cish* and *Socs2* - to determine if there is increased *Stat5b* activity in these leukemias. We saw increased expression of both *Cish* and *Socs2* in leukemic cells from both *Pax5*^+/-^ *x Ebf1*^+/-^ and *Pax5*^+/-^ *x Ebf1*^+/-^ *x Cd79a-Cre x SB* leukemias, which suggests that pStat5 is active in the leukemic cells (Fig. 4H, I). Similar expression results were seen for *Jak1*. We detected a significant increase in *Jak1* mRNA in mice harboring insertions in the *Jak1* locus (Sup. Fig. 4B,C). This increase in *Jak1* transcription significantly increased expression of JAK1 protein in leukemic samples with an SB transposon insertion in the *Jak1* gene locus (Sup. Fig. 4D,E). Our findings are consistent with the high rate of STAT5 activation observed in both human and murine B-ALL[11, 20] and underscore the critical role of JAK/STAT5 signaling in B cell leukemia – particularly those with reduced expression of *Pax5* and *Ebf1*.

**Figure 4.**
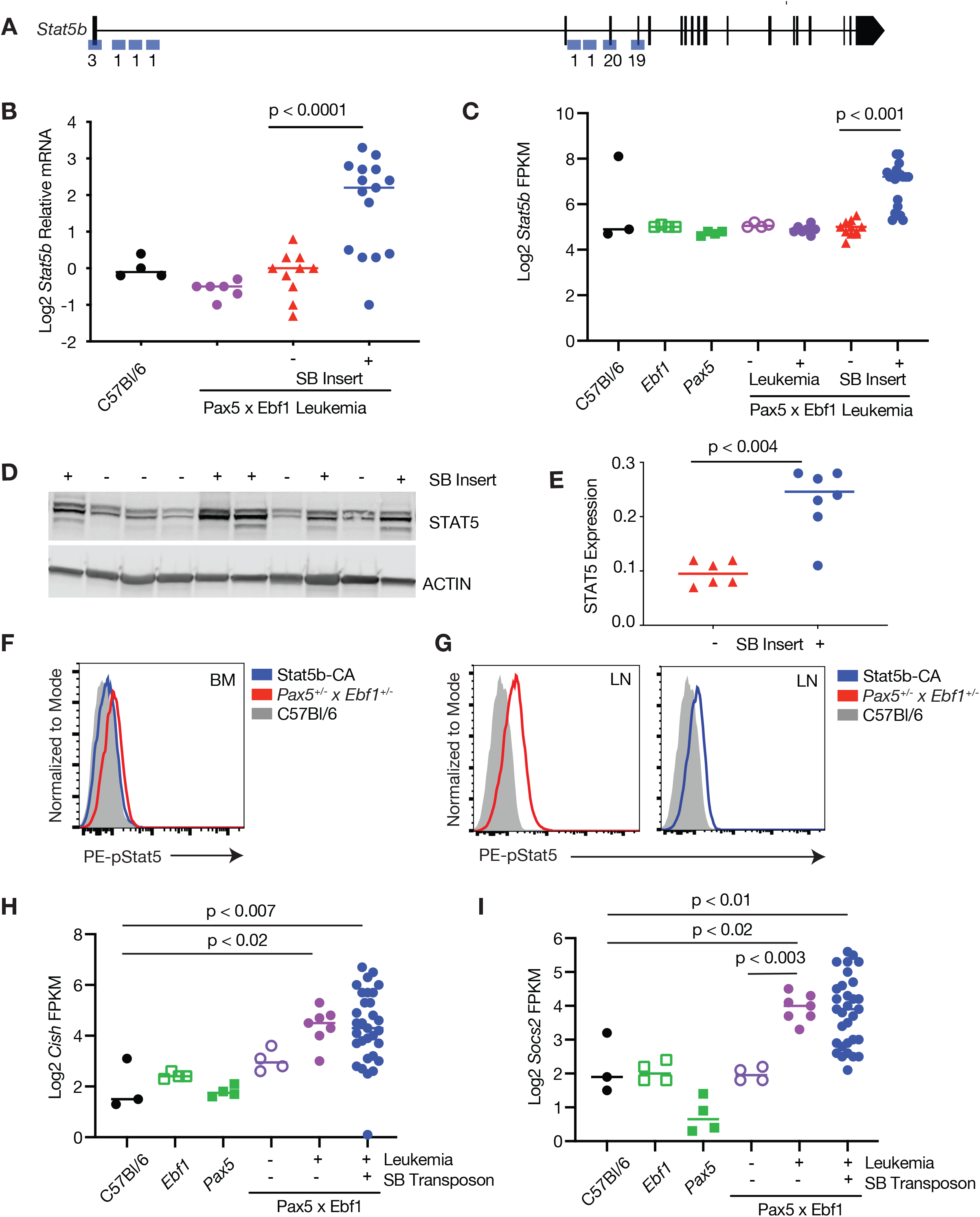
Increased Expression of Stat5b in leukemia. **A** Map of common insertion sites in the *Stat5b* gene; numbers refer to number of insertions at a particular site. **B** Quantitative Real Time PCR (qRT-PCR) for *Stat5b* normalized to *Actin* in progenitor B cells isolated from the bone marrow of WT (black, n=4) mice, and leukemic cells isolated from the lymph nodes of *Pax5*^+/-^ *x Ebf1*^+/-^ (purple, n=6) and SB *Pax5*^+/-^*x Ebf1*^+/-^ mice. The samples from the SB *Pax5*^+/-^*x Ebf1*^+/-^ mice were split between those with (blue, n=15) or without (red, n=10) an insertion in the *Stat5b* locus. The normalized values were log2 transformed and an ordinary one-way ANOVA with Holm-Sidak’s multiple comparison test was used to determine significance. The line represents the median value. **C** Log2 transformed fragments per kilobase of exon model per million reads mapped (FPKM) values from WT (black filled, n=3), *Pax5*^+/-^(green filled, n=4), *Ebf1*^+/-^(green open, n=4), *Pax5*^+/-^ *x Ebf1*^+/-^ pre-leukemic (purple open, n=4), *Pax5*^+/-^ *x Ebf1*^+/-^ leukemic (purple filled, n=7), and SB *Pax5*^+/-^ *x Ebf1*^+/-^ leukemic samples with (blue filled, n=20) or without (red filled, n=11) a transposon insertion in *Stat5b* locus. A Kruskal-Wallis test with Dunn’s multiple comparison test was used to test for significance. The line represents the median value. **D** Western blot analysis showing increased expression of STAT5. The + or - indicates the presence or absence of a SB transposon insertion in each representative sample. **E** Plotted ratio of STAT5 to actin from the western blot. Samples were plotted according to transposon insert status where those samples without a transposon insert are red (n=7) and those with a transposon insert are blue (n=6). Significance was determined using an unpaired student t-test and the line represents the median. **F** Flow cytometric analysis of bone marrow cells from *Pax5*^+/-^ *x Ebf1*^+/-^ leukemic mice. Representative flow cytometric analysis of pSTAT5 expression in cells where doublets were gated out, a lymphocyte gate was applied, and cells were further gated on B220 and AA4.1. This is representative of 5 independent experiments. **G** Flow cytometric analysis of leukemic B cells from *Pax5*^+/-^ *x Ebf1*^+/-^ leukemic mice. Lymph node cells from leukemic mice or bone marrow cells from WT mice were activated with IL-7 for 30 minutes and subjected to flow cytometric analysis for pSTAT5 expression. Doublets were gated out, a lymphocyte gate was applied, and cells were further gated on B220 and AA4.1. This is a representative plot of 4 independent experiments. **H** Log2 transformed FPKM values for *Cish* from WT (black filled, n=3), *Pax5*^+/-^(green filled, n=4), *Ebf1*^+/-^(green open, n=4), *Pax5*^+/-^ *x Ebf1*^+/-^ pre-leukemic (purple open, n=4), *Pax5*^+/-^ *x Ebf1*^+/-^ leukemic (purple filled, n=7), and SB *Pax5*^+/-^ *x Ebf1*^+/-^ leukemic (blue filled, n=31) samples. An ordinary one-way ANOVA with multiple comparisons was used to test for significance. The line represents the median value. **I** Log2 transformed FPKM values for *Socs2* from WT (black filled, n=3), *Pax5*^+/-^(green open, n=4), *Ebf1*^+/-^(green filled, n=4), *Pax5*^+/-^ *x Ebf1*^+/-^ pre-leukemic (purple open, n=4), *Pax5*^+/-^ *x Ebf1*^+/-^ leukemic (purple filled, n=7), and SB *Pax5*^+/-^ *x Ebf1*^+/-^ leukemic (blue filled, n=31) samples. An ordinary one-way ANOVA with Holm-Sidak’s test for multiple comparisons was used to test for significance. The line represents the median value.

### Loss of *Cblb* cooperates with reduced expression of *Pax5* and *Ebf1* to more rapidly induce leukemia

The other top hit in our mutagenesis screen was *Cblb*, which was targeted in almost 2/3 of our leukemias. Transposon insertional analysis from RNA-seq suggested that *Cblb* expression would be reduced as the majority of the SB gene fusions detected involved splicing in exons 6-9 (Fig. 5A). Consistent with this idea, both spontaneous *Pax5*^+/-^ *x Ebf1*^+/-^ leukemias, and SB-induced *Pax5*^+/-^ *x Ebf1*^+/-^ leukemias with an SB insertion in *Cblb*, showed significantly reduced *Cblb* mRNA expression (4.6-fold) compared to WT controls (Fig. 5B). Similar results were seen for CBLB protein expression as SB-induced leukemias with an SB insertion in the *Cblb* gene exhibited significantly lower expression of CBLB protein (1.8-fold) than SB-induced leukemias without an insert (Fig. 5C,D). To determine the role of reduced *Cblb* expression in leukemic transformation, we crossed our *Pax5*^+/-^ *x Ebf1*^+/-^ mice to *Cblb*^*-/-*^ mice. *Pax5*^+/-^ *x Ebf1*^+/-^ *x Cblb*^*-/-*^ mice developed B-ALL and died significantly faster than *Pax5*^+/-^ *x Ebf1*^+/-^ mice, demonstrating that *Cblb* acts as a tumor suppressor in progenitor B cells (Fig. 5E).

**Figure 5.**
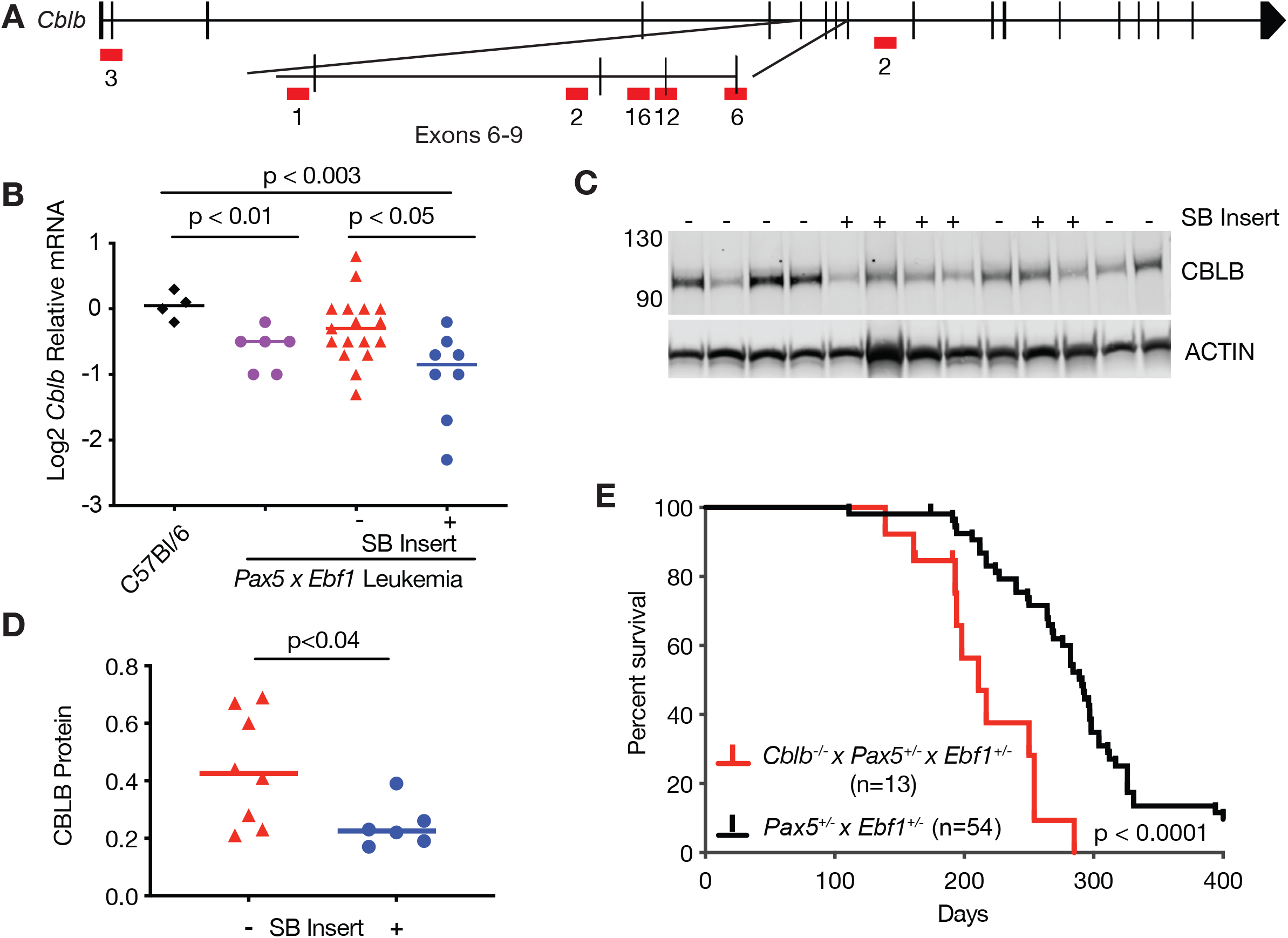
Loss of *Cblb* accelerates the onset of B cell ALL. **A** Map of common insertion sites in the *Cblb* gene; numbers represent number of insertions at a specific site. **B** qRT-PCR for *Cblb* normalized to *Actin* in progenitor B cells isolated from the bone marrow of WT (black, n=4) mice, and leukemic cells isolated from the lymph nodes of *Pax5*^+/-^ *x Ebf1*^+/-^ (purple, n=6) and SB *Pax5*^+/-^*x Ebf1*^+/-^ mice. The samples from the SB *Pax5*^+/-^*x Ebf1*^+/-^ mice were split between those with (blue, n=8) or without (red, n=17) an insertion in the *Cblb* locus. The normalized values were log2 transformed and significance was determined using a Kruskal-Wallis test with Dunn’s multiple comparison test. The line represents the median value. **C** Western blot analysis showing expression of CBLB. The + or - indicates the presence or absence of a SB transposon insertion in each representative sample. **D** Plotted ratio of CBLB to actin from the western blot. Samples were plotted according to transposon insert status where those samples without a transposon insert are red (n=6) and those with a transposon insert are blue (n=8). Significance was determined using an unpaired student t-test and the line represents the median. **E** Kaplan-Meier survival analysis of mice comparing *Pax5*^+/-^ *x Ebf1*^+/-^ leukemic mice (n=51) and *Cblb*^*-/-*^ *x Pax5*^+/-^ *x Ebf1*^+/-^ (n=13) leukemic mice.

### Reduced levels of *Myb* cooperate with *Pax5* and *Ebf1* heterozygosity to more rapidly induce leukemia

*Myb* was another frequent target of our mutagenesis screen. SB transposon insertions were scattered throughout the *Myb* gene locus, suggesting that this would result in decreased expression of *Myb* (Fig. 6A). Spontaneous *Pax5*^+/-^ *x Ebf1*^+/-^ leukemias showed a clear trend towards reduced *Myb* expression. Consistent with this observation, in SB-induced *Pax5*^+/-^ *x Ebf1*^+/-^ leukemias we saw a decrease in *Myb* expression in leukemias that lacked an SB insertion in *Myb* (1.5-fold, Fig. 6B) and an additional significant decrease in leukemias with an SB insertion in the Myb (2.3-fold, Fig. 6C). Thus, downregulation of *Myb* appears to be a general feature of *Pax5*^+/-^*x Ebf1*^+/-^ leukemias. In SB-induced *Pax5*^+/-^ *x Ebf1*^+/-^ leukemias with insertions in the *Myb* gene, there was also a significant reduction (2.8 fold, Fig. 6D, E) at the protein level. Importantly, we found that *Myb* expression as assessed by RNA-Seq correlated with age of death - leukemias with less *Myb* were more aggressive resulting in earlier lethality (Fig. 6F).

**Figure 6.**
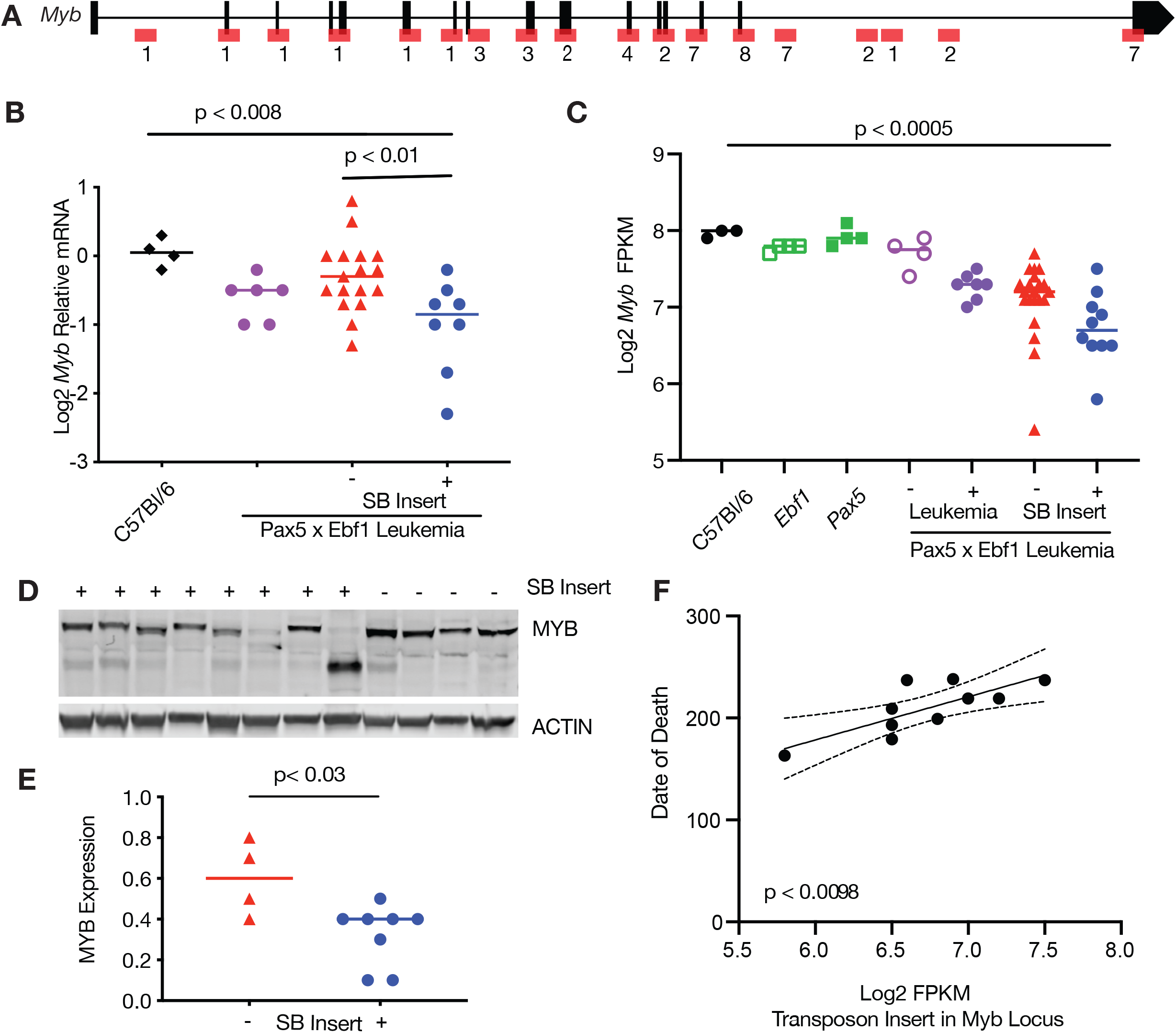
Loss of MYB expression results in worse outcome in ALL. **A** Map of common insertion sites in the *Myb* gene; numbers refer to number of insertions at a specific site. **B** qRT-PCR for *Myb* normalized to *Actin* in progenitor B cells isolated from the bone marrow of WT (black, n=4) mice, and leukemic cells isolated from the lymph nodes of *Pax5*^+/-^ *x Ebf1*^+/-^ (purple, n=6) and SB *Pax5*^+/-^*x Ebf1*^+/-^ mice. The samples from the SB *Pax5*^+/-^*x Ebf1*^+/-^ mice were split between those with (blue, n=8) or without (red, n=17) an insertion in the *Myb* locus. The significance was tested using an ordinary one-way ANOVA with Holm-Sidak’s multiple comparison test and the lines represent median. **C** Log2 transformed fragments per kilobase of exon model per million reads mapped (FPKM) values from WT (black filled, n=3), *Pax5*^+/-^ (green filled, n=4), *Ebf1*^+/-^(green open, n=4), *Pax5*^+/-^ *x Ebf1*^+/-^ pre-leukemic (purple open, n=4), *Pax5*+/-*x Ebf1*^+/-^ leukemic (purple filled, n=7), and SB *Pax5*^+/-^ *x Ebf1*^+/-^ leukemic samples with (blue filled, n=10) or without (red filled, n=21) a transposon insertion. Significance was tested using a Kruskal-Wallis test with Dunn’s multiple comparison test and the line represents median. **D** Western blot analysis showing decreased expression of MYB in SB leukemia samples harboring a transposon insertion. The + or - indicates the presence or absence of a SB transposon insertion in each representative sample. **E** Plotted ratio of MYB to actin from the western blot in panel d. Samples were plotted according to transposon insert status where those samples without a transposon insert are red (n=4) and those with a transposon insert are blue (n=8). Significance was determined using an unpaired student t-test and the line represents the median. **F** Linear regression analysis comparing date of death versus FPKM value for leukemic samples harboring a transposon insertion. The dashed lines represent 95% confidence bands.

### PDK1-signaling pathway is deregulated in *Pax5*^+/-^ *x Ebf1*^+/-^ leukemias

In addition, to gene alterations directly regulated by SB transposition, we also noted a number of genes whose expression was significantly altered in *Pax5*^+/-^ *x Ebf1*^+/-^ leukemias relative to non-transformed controls. These included genes such as the tumor suppressor *Bach2*, which was significantly reduced (Sup. Fig. 5A). Intriguingly, we also noted dramatic (5.7 fold) downregulation of Asparagine synthetase in these leukemias, which may explain their susceptibility to L-Asparaginse[21](Sup. Fig. 5B). Other genes were strikingly upregulated including *Pdk1* (3.0-fold) and its downstream targets *Sgk3*, and *Rhebl1* (Fig. 7A,B,C). Conversely *Tsc2*, which inhibits this pathway, was downregulated (Sup. Fig. 5C,D); this pathway has been previously shown to enhance mTORC1 function and ultimately MYC expression[22, 23]. To determine whether PDK1 plays a critical role in maintaining viability of *Pax5*^+/-^ *x Ebf1*^+/-^ leukemias, we treated two cell lines generated from *Pax5*^+/-^ *x Ebf1*^+/-^ leukemias in vitro with either vehicle control or the PDK1 inhibitor (GSK2334750). Treatment of these cell lines with the PDK1 inhibitor resulted in a dose-dependent decrease in the survival of these cell lines suggesting that this might be a useful treatment for B-ALL subsets with reduced *Pax5* and *Ebf1* expression (Fig. 7D).

**Figure 7.**
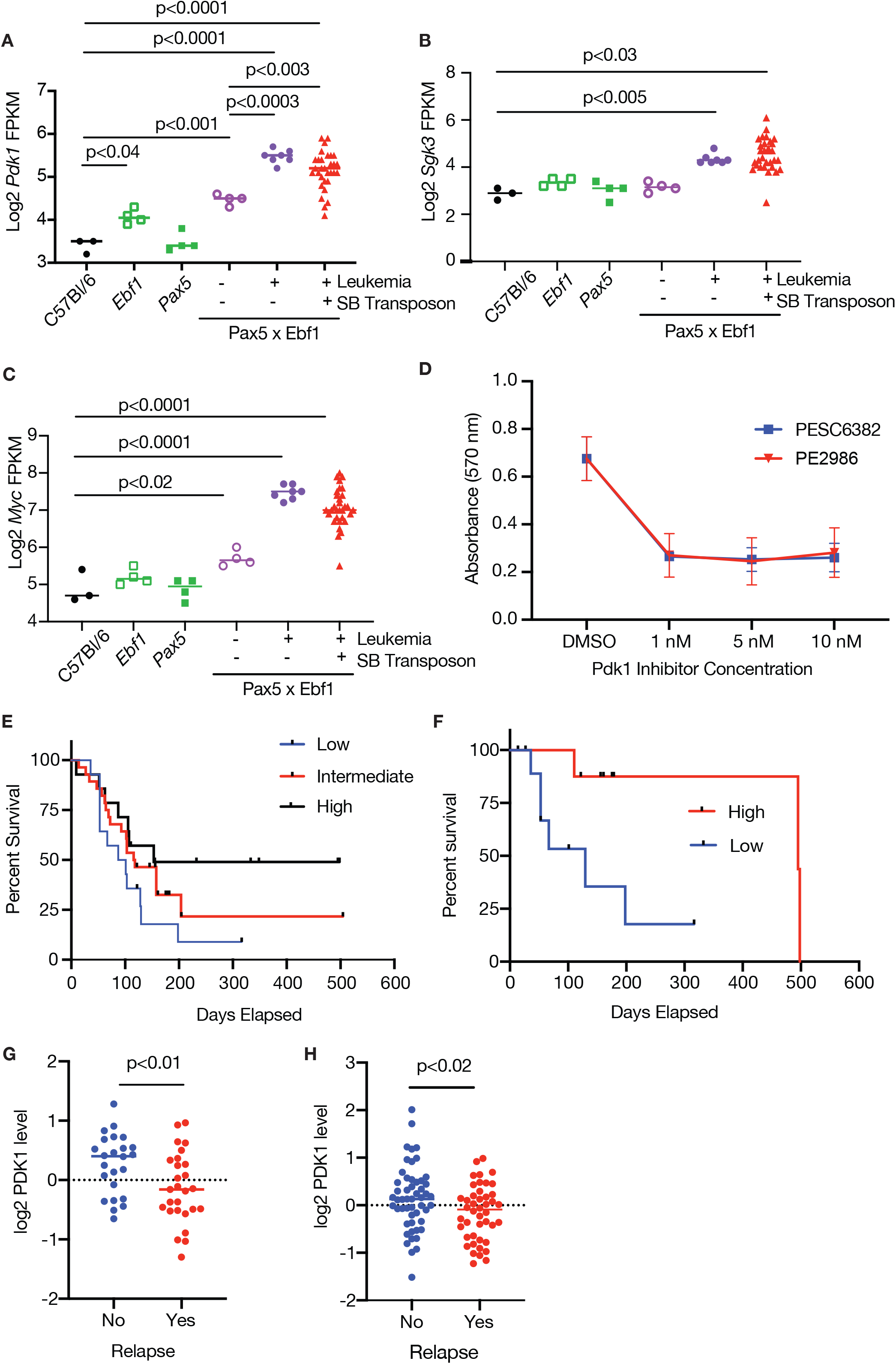
Inhibition of PDK1 blocks leukemic proliferation. **A** Log2 transformed FPKM values for *Pdk1* from the RNA-seq datasets. Significance was determined by an ordinary one-way ANOVA with Holm-Sidak’s multiple comparison test. **B** Log2 transformed FPKM values for *Sgk3* from the RNA-seq datasets. Significance was tested using a Kruskal-Wallis test with Dunn’s multiple comparison test. **C** Log2 transformed FPKM values for *Myc* from the RNA-seq datasets. Significance was determined by an ordinary one-way ANOVA with Holm-Sidak’s multiple comparison test. **D** PDK1 inhibitor blocks growth. MTT assay was performed on two *Pax5*^+/-^ *x Ebf1*^+/-^ leukemic cell lines generated from lymph node cells from leukemic mice. The cells were subjected to differing concentrations of GSK2334470. Each dot represents the average of two biological replicates - each biological replicate represents the mean of triplicate technical replicates within an experiment. Error bars represent the range between experiments. **E** Overall survival of *Bcr-Abl* patients. *Bcr-Abl* patients were separated into three equal groups representing higher (black line, n=14), intermediate (red line, n=28) and lower (blue line, n=14) levels of PDK1. Patients with lower levels of PDK1 did significantly worse than patients with higher PDK1 (p=0.04, Log Rank test for trend). **F** Overall survival of *Bcr-Abl* young adult patients. Young adult patients were separated into two equal groups representing higher (red line, n=9) and lower (blue line, n=9) levels of PDK1. Patients with lower levels of PDK1 did significantly worse than patients with higher PDK1 (p=0.02, Log Rank (Mantel-Cox) Test). **G** PDK1 expression of *Bcr-Abl* patients split by relapse status. *Bcr-Abl* patients were separated into two groups representing no relapse (blue dots, n=14), and relapse (red dots, n=14) levels of PDK1 were graphed. Patients with lower levels of PDK1 did significantly worse than patients with higher PDK1 (p<0.01, unpaired T-test). **H** PDK1 expression of *B-Nos* patients split by relapse status. Patients were separated into two groups representing no relapse (blue dots, n=14), and relapse (red dots, n=14) and levels of PDK1 were graphed. Patients with lower levels of PDK1 did significantly worse than patients with higher PDK1 (p<0.02, unpaired T-test)

To examine a possible role for PDK1 expression in human ALL, we examined ALL patient samples using a reverse phase proteomics approach[24]. PDK1 was expressed in five subsets of B-ALL but expression varied widely (Sup. Fig. 5E). We examined PDK1 expression in the two largest cohorts B-NOS and BCR-ABL+ leukemias. PDK1 levels did not correlate with overall survival in B-NOS patients (data not shown). In contrast, BCR-ABL+ patients with the highest levels of PDK1 expression did significantly better than those with lower PDK1 expression (Fig. 7E). The difference in overall survival appears to be driven most strongly by young adults, as they showed the most dramatic difference in overall survival (Fig. 7F). Finally, we examined PDK1 expression in patients with BCR-ABL and B-NOS leukemia based on relapse status. In both subsets of leukemia, lower levels of PDK1 correlated with relapse (Fig. 7G,H). Thus, PDK1 appears to play an important role in B-ALL survival or proliferation, but patients with the highest level of PDK1 expression respond better to therapy.

### Pax5 x Ebf1 leukemia show common transcriptional variation patterns across mouse and human

To determine if the murine leukemias that developed in our sleeping beauty screen are similar to subsets of human B-ALL, we quantified inter-leukemia transcriptional variation using our newly developed gene <u>c</u>luster <u>e</u>xpression <u>s</u>ummary <u>s</u>core (GCESS)[25]. Using this approach, we first examined inter-leukemia transcriptional variation in distinct human leukemia datasets. There was notable heterogeneity between human B-ALL subsets. However, we could identify clusters with similar variations in gene expression (Fig. 8A). We used the GCESS approach to establish transcriptional similarity between these human leukemias and our murine sleeping beauty transposon induced leukemias. As shown in figure 8B, there were two distinct transcriptional variants in our SB dataset. One of these SB leukemia subsets showed a similar gene expression signature to one of the conserved human leukemia subsets (fishers exact test, p=4.3e-07). Thus, the leukemias that developed in our SB system are similar to human leukemias, thereby validating our approach as a useful model of human leukemia.

**Figure 8.**
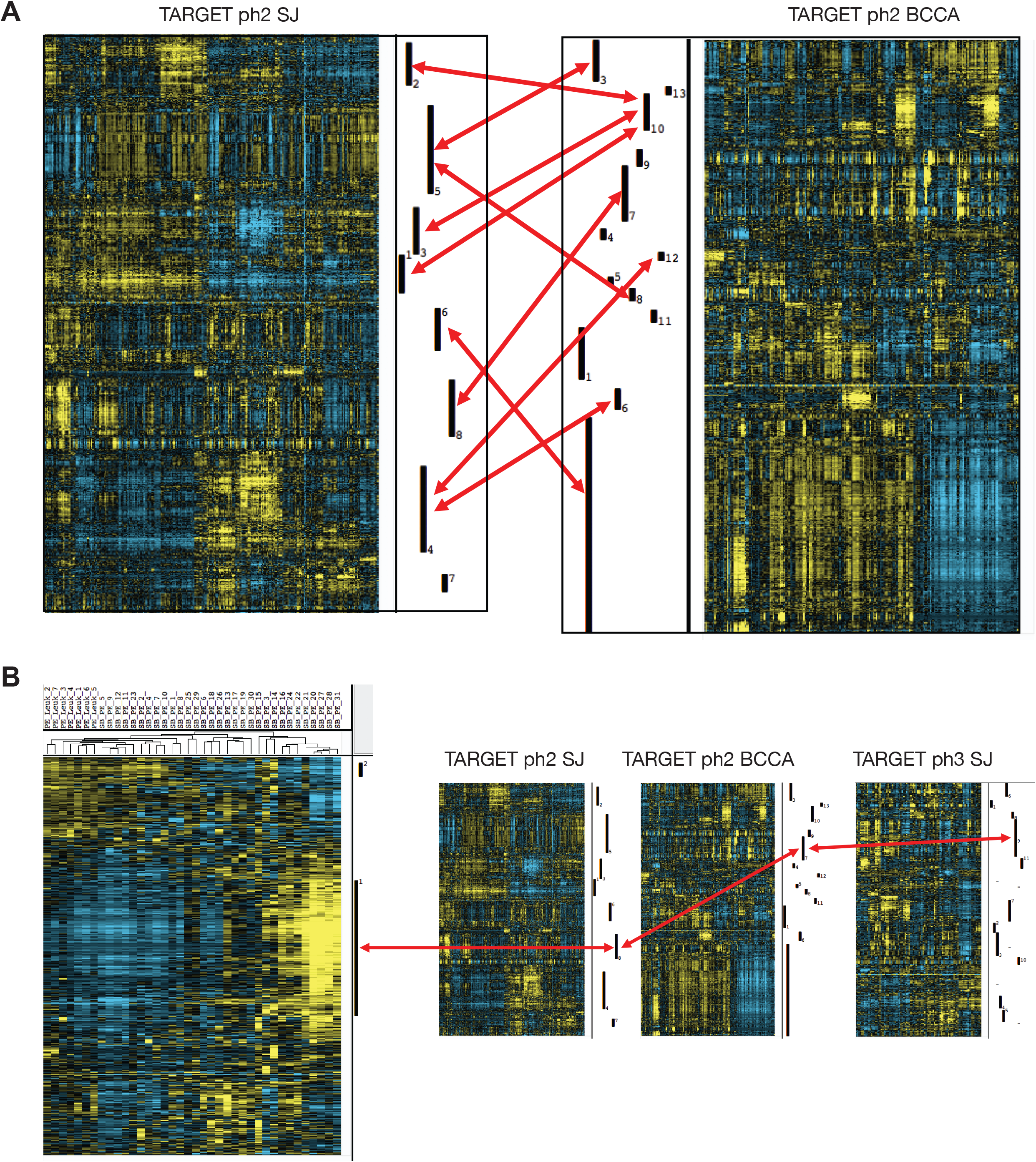
Transcriptome profiles from leukemic progenitor B cells show common interleukemic transcriptional variation across human and mouse samples. **A** Common transcriptional patterns identified in RNA-Seq from human leukemias. Top ∼8000 gene values defined by SD in ph2_SJ and ph2_BCCA separately identify conserved transcriptional patterns to be present. Red bars indicate highly significantly enriched sets of conserved genes to be present across Human ALL datasets via Fisher exact test comparison of gene cluster membership. Gene transcript values derived from human leukemias were log transformed and mean centered within each species. Invariant (low SD) genes were removed prior to unsupervised average linkage clustering. Conservation was apparent despite the fact that the SJ set was summarized as gene symbols while the BCCA set was summarized to ENSEMBL ids. Transcripts with increased levels are shown in yellow, while transcripts with decreased levels are shown in blue. **B** Gene cluster overlap analyses comparing clusters derived from human and mouse tumors show that the variation present in our mouse dataset represents one of the clear variations present and conserved in all of the human samples. Gene lists for the conserved cluster from each dataset are provided in Supplementary Table S2.

To better characterize these leukemias, we utilized ENRICHR to examine gene lists from the GCESS of each of the datasets from the conserved murine and human leukemias (Sup. Table 1)[26, 27]. Consistent with other findings in this study, we found upregulation of cytokines and cytokine receptor genes, as well as genes involved in JAK/STAT5 signaling. In addition, NFκB signaling was significantly altered, which is consistent with work from multiple groups on NFκB in B cell development and leukemia [20, 28, 29]. A surprising observation was a strong myeloid gene signature in the human and murine leukemias. There are two potential reasons for this. First, these leukemias could be infiltrated with myeloid cells. Alternatively, the leukemias could have lost lineage fidelity and begun to express myeloid genes. Since PAX5, EBF1 and IKZF1 all play key roles in enforcing B cell lineage fidelity, and our murine B cell leukemias were relatively pure populations of leukemic blasts, we favor this later possibility. Thus, subsets of *Pax5*^+/-^ *x Ebf1*^+/-^ leukemia exhibit some myeloid characteristics.

## Discussion

Genes encoding the transcription factors *PAX5, EBF1* and *IKZF1* frequently exhibit loss of one allele or express loss-of-function mutations in human B cell leukemia[2, 4, 30]. However, their role in B cell transformation is not entirely clear. It is likely that loss of function mutations in these transcription factors affect B cell differentiation. Previous studies using inducible Pax5 mutants in murine models of B cell leukemia suggest that this plays a role[31]. However, *Pax5*^+/-^ and *Ebf1*^+/-^ mice do not develop leukemia, while *Ikzf1*^+/-^ mice have only been shown to develop T cell leukemia. This raises the question of how mutations in these genes promote transformation. We previously demonstrated that combining loss of one allele of either *Pax5* or *Ebf1* with a constitutively active *Stat5* transgene (referred to as *Stat5b-CA*) led to rapid onset of B cell leukemia. A key feature of these leukemias is that *Ebf1* expression was reduced ∼50% in *Stat5b-CA x Pax5*^+/-^ leukemias, while *Pax5* expression was comparably reduced in *Stat5b-CA x Ebf1*^+/-^ leukemias[11]. These findings suggested that perhaps compound haploinsufficiency for these transcription factors might be key for promoting transformation. Herein we demonstrated that this is the case as *Pax5*^+/-^ *x Ebf1*^+/-^ mice develop B cell leukemia with high penetrance. Importantly, this was not a phenomenon restricted to this pair of transcription factors as we saw qualitatively similar onset of B cell leukemia in *Pax5*^+/-^ *x Ikzf1*^+/-^ and *Ebf1*^+/-^ *x Ikzf1*^+/-^ mice. Nor was this observation restricted to B cell leukemia as we observed that *Tcf7*^+/-^ *x Ikzf1*^+/-^ mice gave rise to highly penetrant T cell leukemia. Thus, compound haploinsufficiency for transcription factors that play key roles in either B cell or T cell development can promote transformation.

The mechanism by which compound haploinsufficiency promotes transformation remains unclear. Previous studies suggested that this may be due to defective DNA repair upon reduced *Ebf1* expression. We were unable to validate defects in *Rad51, Rad51ap* expression or increased γH2AX expression in pre-B cells from Ebf1^+/-^ mice. However, we did observe that a subset of human leukemias expressed a hypermutated phenotype and that leukemias with *CDKN2A, EBF1* and *IKZF1* missense mutations were enriched in this subset. It is possible that the newly described BCR-ABL-like subset of B-ALL (which is enriched in leukemias with mutations in *CDKN2A, IKZF1* and *EBF1*) might be characterized by the hypermutated phenotype. This should be examined further and if confirmed suggests that these leukemias may be more susceptible to immunotherapy-based treatments.

An alternative mechanism by which compound haploinsufficiency promotes transformation could involve loss of lineage fidelity. Consistent with this hypothesis, EBF1 and PAX5 have both been shown to play important roles in restricting cells to the B cell lineage. Consistent with this observation, B cell progenitors in *Pax5*^+/-^ *x Ebf1*^+/-^ mice retain T cell lineage potential[32]. Moreover, we found that compound haploinsufficiency for *Pax5* and *Ebf1* promotes increased penetrance of B cell and T cell leukemia on an *Ikzf1*^+/-^ background. Thus, decreased expression of transcription factors that are only expressed in B cells (PAX5 and EBF1) paradoxically enhance the development of T cell leukemia in *Ikzf1*^+/-^ mice. Although the mechanism remains unclear, it is possible that in some cases T-ALL may emerge from a B cell progenitor. This may have implications for how such leukemias develop resistance to therapy if the key progenitor is more closely linked to B cell rather than T cell development. Finally, our finding that a subset of murine *Pax5*^+/-^ *x Ebf1*^+/-^ leukemias, as well as their similar human B-ALL counterparts, exhibit a strong myeloid gene signature also suggests that loss of lineage fidelity may be a key feature of this disease. Alternatively, the myeloid signature could arise due to preferential infiltration of this type of leukemia with myeloid cells. This is certainly a possibility, especially for leukemias in the human datasets. However, our murine leukemias are strongly enriched for B lineage cells. Thus, we favor an explanation in which the myeloid gene signature arises due to aberrant expression of myeloid genes in B cell leukemic blasts.

To gain a better understanding of the molecular alterations that cooperate with *Pax5* and *Ebf1* haploinsufficiency to promote transformation we carried out an unbiased SB transposon screen in *Pax5*^+/-^ *x Ebf1*^+/-^ mice. These studies identified two major pathways that cooperate with *Pax5* and *Ebf1* haploinsufficiency to drive transformation. First, we found gain-of-function mutations for *Jak1* or *Stat5b* in almost all of our leukemias in this screen. This finding underscores in an unbiased way the critical role of JAK/STAT5 signaling in B cell transformation. In a previous SB mutagenesis study targeting the STAT5 pathway, we were able to induce more rapid leukemia onset than SB mice with only *Pax5*^+/-^ *x Ebf1*^+/-^ predisposing mutations (average onset of leukemia ∼72 versus 302 days, respectively) ^*[33]*^. This suggests that changes needed to activate STAT5 may take longer to arise than secondary loss-of-function mutations to *Pax5, Ebf1*, or other transcription factors.

The second major pathway identified involves CBL-B, and to a much lesser extent the related family member CBL. The mechanism by which *Cblb* loss-of-function affects transformation is unclear. However, the fact that these are loss-of-function mutations is supported by the observation that crossing *Cblb*-deficiency onto the *Pax5*^+/-^ *x Ebf1*^+/-^ background accelerated the onset of leukemia and increased overall penetrance. Our SB screen suggest that *CBLB* mutations should be examined in greater detail in human B-ALL.

A number of other target genes were identified in our SB screen, although none were as prevalent as the mutations in *Jak1, Stat5b* or *Cblb*. These include SB insertions in several cytokine/receptor genes that have previously been shown to be involved in transformation including *Il2rb, Gh, Csf2*, as well as the histone acetyltransferase *Ep300*. Finally, we noted relatively frequent mutations in the transcription factor *Myb*. The finding that *Myb* was targeted by SB in our leukemias was initially not surprising as *Myb* has previously been identified as an oncogene. However, what was surprising is that the mutations in *Myb* were <u>loss-of-function</u> mutations resulting in <u>reduced</u> *Myb* expression. This leads to the somewhat surprising observation that *Myb* acts operationally as a tumor suppressor in this context and parallels our previous observation that NFκB also acts as a functional tumor suppressor in progenitor B cells[20, 34]. Since MYB plays a role in B cell differentiation[35] this likely reflects a role for MYB in blocking differentiation at the highly proliferative pre-B cell stage.

In addition to targets directly identified by SB integration, our RNA-Seq analysis of *Pax5*^+/-^ *x Ebf1*^+/-^ and *SB x Pax5*^+/-^ *x Ebf1*^+/-^ leukemias also identified other deregulated signaling pathways. One of the most prominently deregulated pathways involved the serine/threonine kinase PDK1. PDK1 expression was modestly but significantly upregulated in *Ebf1*^+/-^ and *Pax5*^+/-^ *x Ebf1*^+/-^ pre-leukemic progenitor B cells, and significantly further elevated in *Pax5*^+/-^ *x Ebf1*^+/-^ leukemias. The mechanism by which increased PDK1 expression promotes transformation is unclear. However, previous studies have shown that PDK1 interacts with SGK1/3 to inhibit TSC2 function and expression[22]. This results in increased function of RHEB or RHEBL1, which in turn promote mTOR function and ultimately MYC expression[22]. Consistent with this model we found that *Tsc2* expression levels were reduced in *Pax5*^+/-^ *x Ebf1*^+/-^ leukemias while *Sgk3, Rhebl1* and *Myc* levels were increased. An alternative pathway that could also be affected by PDK1 involves PDK1-dependent activation of PLK1, which in turn has been shown to phosphorylate and activates MYC[23]. Thus, there are a number of potential mechanisms by which increased PDK1 expression could promote transformation. What is clear is that PDK1 inhibitors effectively blocked proliferation of *Pax5*^+/-^ *x Ebf1*^+/-^ primary leukemia cell lines in vitro. Since there are currently a number of PDK1 inhibitors available with some demonstrating efficacy in preclinical trials[36, 37], our findings suggest that PDK1 inhibition might be an effective strategy for treating B cell leukemias that exhibit reduced expression of *Pax5* and *Ebf1*.

## ACKNOWLEDGEMENTS

We thank A. Rost, for technical assistance with mouse breeding; the University of Minnesota’s Supercomputing Institute for providing computing and bioinformatic resources; Dr. Meinrad Busslinger (*Pax5*^+/-^), Dr. Rudolf Grosschedl (*Ebf1*^*-/-*^), Dr. Peter Johnson (*Cebpa*^*-/-*^), Dr. Andrew Wells (*Ikzf1*^*-/-*^) and Dr. David Largaespada (Rosa26^LSL-SB11^ *T2/OncxRosa26*^*LSL-SB11*^) for providing the indicated mouse strains. The results published here are in part based upon data generated by the Therapeutically Applicable Research to Generate Effective Treatments (TARGET) initiative, phs000218, managed by the NCI. This work was supported by a Cancer Research Institute Investigator award, a Leukemia and Lymphoma Society Scholar award, funding from the UMN Masonic Cancer center and grants from the NIH (RO1 CA232317) to MAF. TKS was supported by grants from the Randy Shaver Cancer Research and Community Fund, NIH NCI (R21 CA216652), and the Masonic Cancer Center. ALS was supported by NCI (CA211249) and Masonic Cancer Center Support Grant (CA077598). SMK was supported by CPRIT MIRA RP 160693 and NIH/NCI P50 CA100632-09.

## Authors’ Contribution

### Conception and design

L. Heltemes-Harris and M. Farrar

### Development of Methodology

L. Heltemes-Harris, A. Sarver, S. Kornblau and M. Farrar

### Acquisition of data (provided animals, acquired and managed patients, provided facilities, etc.)

L. Heltemes-Harris, G. Hubbard, A. Sarver, T. Knutson, S. Kornblau and M. Farrar

#### Analysis and interpretation of data (e.g., statistical analysis, biostatistics, computational analysis)

L. Heltemes-Harris, G. Hubbard, R. LaRue, T. Starr, S. Munro, T. Knutson, C. Henzler, A. R. Yang, A. Sarver, S. Kornblau and M. Farrar.

### Writing, review, and/or revision of the manuscript

L. Heltemes-Harris, G. Hubbard, R. LaRue, T. Starr, S. Munro, C. Henzler, R. Yang, A. Sarver, S. Kornblau and M. Farrar.

#### Administrative, technical, or material support (i.e., reporting or organizing data, constructing databases)

L. Heltemes-Harris

### Study Supervision

L. Heltemes-Harris, M. Farrar

