## Supplemental methods and figures for "Identification of mutations that cooperate with defects in B cell transcription factors to initiate leukemia"

**Supplementary Data**

**Sleeping Beauty transposon screen identifies mutations that cooperate with defects in B cell transcription factors to initiate leukemia**

*Heltemes-Harris et al.,*

#### Supplemental materials and methods

##### Cells

Purification of progenitor B cells was done using a two-step process. Cells were labeled with FITC-labeled α-Igλ (Southern Biotech), α-Igκ (Southern Biotech), α-Ly6G (GR-1, eBioscience) and α-Ter119 (eBioscience) followed by labeling with anti-FITC microbeads (Miltenyi) to deplete mature and non-B cells. Cells were labeled with anti-CD19 microbeads (Miltenyi) and cells bound to the column were collected and used in further experiments.

*Pax5*$^{+/-}$ *x Ebf1*$^{+/-}$ leukemic cell lines (called PE2986 and PE6382) were generated by plating leukemic cells isolated from the lymph nodes of the sick animals on OP9 feeder layers supplemented with recombinant mouse IL7 (Tonbo Biosciences). Once the cells were growing, cells were removed from the feeder layers and grown in suspension on recombinant mouse IL7 (Tonbo Biosciences).

##### Sleeping Beauty Mutagenesis

*Pax5*$^{+/-}$ and *Ebf1*$^{+/-}$ mice were crossed to *Cd79a-Cre* mice, which express CRE recombinase in developing and mature B cells. *Pax5*$^{+/-}$*xEbf1*$^{+/-}$*xCd79a-cre* mice were then further crossed to mice that harbor a concatemer of mutagenic transposon vectors

(T2/Onc) on chromosome 1 or 15 as well as  a Cre-inducible SB transposase gene (Rosa26$^{LSL-SB11}$) (*T2/OncxRosa26$^{LSL-SB11}$* combination, referred to as *SB* herein) (1). The Rosa26$^{LSL-SB11}$ transposase transgene also contains a *Gfp* cDNA that is removed upon CRE-mediated recombination and therefore allows identification of cells in which CRE recombinase is active and the *SB11* transposase enzyme is expressed. We generated 34 leukemic mice with 12 mice from the Chromosome 1 system and 19 mice from the Chromosome 15 system.

**Flow Cytometry**

Single cell suspensions were prepared from the bone marrow, lymph nodes and spleen and stained with antibodies listed in table below and run on a LSRII flow cytometer (BD Biosciences); data was analyzed using FlowJo software (Treestar).

| **Antibody - Flow Cytometry** | **Company** |
|---|---|
| Anti-IgM | Jackson ImmunoResearch |
| Anti-IgD (11-26) | eBioscience |
| Anti-Igλ | Southern Biotech |
| Anti-Igκ | Southern Biotech |
| Anti-Ter119 | eBioscience |
| Anti-Ly6G (GR-1) | eBioscience |
| Anti-Ly6A/E (Sca1) | Biolegend |
| Anti-CD3 (145-2C11) | Tonbo Biosciences |
| Anti-CD4(RM4-5) | Tonbo Biosciences |
| Anti-CD8a (Ly2) | eBiosciences |
| Anti-CD19 (1D3) | BD Biosciences |
| Anti-CD24 (M1/69) | BD Biosciences |
| Anti-CD25 (PC61.5) | BD Biosciences |
| Anti-CD43 (S7) | eBioscience |
| Anti-CD45R (RA3-6B2) | BD Biosciences |
| Anti-CD90.2 (Thy1.2, 30-H-12) | Tonbo Biosciences |
| Anti-CD93 (AA4.1) | eBioscience |

| | |
|---|---|
| α-CD117 (2B8), | eBioscience |
| α-CD127 (A7R34) | Biolegend |
| α-CD135 (A2F10) | eBioscience |
| α-CD249 (BP1-Ly51) | BD Biosciences |
| α-pre-BCR (SL165) | BD Biosciences |

**RNA-Seq Analysis**

Fragments per kilobase of exon model per million reads mapped (FPKM) values generated from RNA-Seq values were calculated as previously described(2). A value of 0.1 was added to each FPKM estimate, the data was mean centered and log base 2 transformed. 2859 genes were identified with **SD** greater than 1.

**Transposon Insertion Analysis**

Fusions can be identified in RNA-Seq data from SB accelerated tumors that represent the oncogenic transcripts generated from Transposon insertion(2, 3). We identified these fusions using a systematic approach based on identification of RNA sequence that contain a junction between genomic and transposon derived RNA as well as paired reads that span either side of a junction as previously described(3).

**Variant calling on RNA-seq data**

Reads were aligned to the mouse genome (mm10) using BWA (0.7.12)(4). Variants were called with freebayes (v. 0.9.21)(5) ignoring multi-nucleotide polymorphisms and complex events. Wild type calls were filtered using vcffilter to keep only those variants with a minimum total depth of 20 reads and a quality score of 20, and remove all variants also found in dbSNP (v142). The three wild type replicates were then filtered to a minimum depth of 1000 reads, which was determined to be the noise threshold, i.e. the minimum read depth at which the differences between the wild type replicates were minimized. To find the unique PE calls, the PE samples were subsequently filtered to keep only calls with a minimum depth of 1000 and a minimum quality score of 20, and to remove any calls found in either dbSNP (v142) or the filtered wild type samples.

**Gene Set Enrichment Analysis**

Reads were aligned with HiSat2(6) to the mouse genome (mm10) with the SB transposon sequence appended. Counts were estimated using the Subread featureCounts tool(7). Differential expression testing on the count data was conducted using the R package edgeR(8). Ranked gene lists for Gene Set Enrichment Analysis (GSEA)(9) were generated using the following two step transformation. First, P-values from the DE test were transformed by -log10, this transformation spreads out the p-value distribution, which initially has a 0 to 1 range. Then the -log10 P-values for each gene are multiplied by either +1 or -1 depending on the sign of the expression fold change for that gene. Positively regulated genes in the comparison will be multiplied by +1 and negatively regulated genes will be multiplied by -1. This ranked gene list was then used for “Preranked” GSEA analysis using default parameters with the GSEA java program (4.0.1). Gene sets related to DNA Repair were selected from MSigDb and were also curated from the work of Prasad and colleagues (9 total sets were tested for enrichment). The Sigvardsson Gene Set is the set of genes nominated for a DNA repair signature in Table S1 of Prasad et al. 2014(10). In addition to analyzing our RNA-Seq data with GSEA we also used the GSEA program recommended settings to analyze

GEO-deposited microarray data from Prasad et al. 2014 and Ashberg et al. 2013(10, 11).

**Western Blot**

Protein was isolated from 50x$10^6$ leukemic cells using RIPA buffer (Thermo Scientific) with HALT protease and phosphatase inhibitors (ThermoScientific) following standard procedures as described previously(12). Briefly, protein concentration was determined using a Bradford Assay (Sigma). The gels were run according to the BioRad Criterian instruction manual and application guide with 50 ug of protein per well. Blots were visualized on a LiCor Odyssey (Li-Cor Biosciences) and analyzed using Image Studio Lite software (Li-Cor Biosciences).

| Antibody - Western Blot | Company |
|---|---|
| phospho-STAT5 (Tyr694) | Cell Signaling |
| STAT5 (3H7) | Cell Signaling |
| JAK1 | Cell Signaling |
| Cblb (C-20) | Santa Cruz |
| Myb (1-1) | Millipore |
| Actin (AC-15) | Sigma |

**qRT PCR Analysis**

RNA was extracted from 50x$10^6$ leukemic or progenitor B cells purified from C57BL/6, *Pax5+/- x Ebf1*$^{+/-,}$ mice using a RNeasy Mini kit (Qiagen) following the recommended procedures. qPCR was done as previously described (12). RNA concentrations were determined in a Nanodrop (ThermoScientific). 250 ng of RNA was converted to cDNA with the ABI High-Capacity cDNA Reverse Transcription Kit (#4368814) according to the manufacturer's instructions. Gene specific primers are listed in table below. PCR was carried out in an ABI 7900HT system in triplicate using the iTaq Universal SYBR Green (Bio-Rad) in 20 ul reactions containing 200 nm of each forward and reverse primer, 2 ul of cDNA diluted mix (~10 ng) and 10 ul of 2X SYBR Master Mix.

| **PrimeTime qPCR Primers, IDT DNA Technologies** | |
|---|---|
| **Gene** | **Reference Number** |
| *Stat5b* | Mm.PT.58.31682119 |
| *Jak1* | Mm.PT.56a.17266971 |
| *Cblb* | Mm.PT.58.41698502 |
| *Myb* | Mm.PT.58.13223048 |
| *Rad51* | Mm.PT.58.42533544 |
| *Rad51AP* | Mm.PT.58.12874053 |
| *Tcf3* | Mm.PT.58.31845661 |
| *gH2AFX* | Mm.PT.58.45969097.g |

**Inhibitor Assay**

The progenitor B cell lines PE2986 and PE6382 generated from our two of our leukemic mice were used in the an MTT assay (Cayman chemicals) as described in the kit. Cells were plated at 50,000 cells per well with differing dilutions of PDK1 inhibitor GSK2334470 (MCE). Samples were read on a Bio-Tek Plate reader (Bio-Tec) at 570 nm.

**GCESS calculation**

The GCESS is defined as the sum of expression values ($log_2$-transformed and mean centered) of all genes in a particular defined cluster for a single sample. The GCESS quantifies transcriptional variation between tumors. It takes many correlated individual transcript data points and condenses them into a single value (13).

**Reverse Phase Protein Assays (phospho-STAT5)**

129 ALL samples were collected and analyzed using reverse phase protein arrays as previously described (14). Patient sample data is provided in Supplemental Table 2. These samples were collected by the University of Texas M.D. Anderson Leukemia sample bank under the IRB approved protocol Lab01-473 and utilized under protocol Lab05-0654.

**Variant Analysis of TARGET ALL-Phase 2 Dataset**

Whole exome sequencing (WES) and copy number variant (CNV) analysis datasets were explored from the TARGET ALL (phase 2) project. Mutation annotation files (MAFs) and gene level CNV files were downloaded from the Genomic Data Commons (GDC) repository (https://portal.gdc.cancer.gov) using the gdc-client (ver. 1.5.0). These datasets were generated by the GDC harmonized analysis pipeline. Primary tumor-normal matched samples were isolated (717 MuTect2 MAFs and 293 CNV files) and T-cell derived samples were removed, leaving 486 B-cell derived samples for analysis. Data tables were imported into R (ver. 3.6.0) and all filtering and plotting was completed using tidyverse package functions (ver. 1.3.0). The MAFs included variant annotations from Variant Effect Predictor (VEP), Polymorphism Phenotyping (PolyPhen-2), and Sorting Intolerant from Tolerant (SIFT) software. These annotations and the copy number data were used to label each sample as "affected" if it contained loss of function variants affecting various genes of interest (EBF1, PAX5, IKZF1, CDKN2A) or contained certain gene-fusions (ETV6-RUNX1, TCF3-PBX1). Samples were assumed to contain loss of function alleles for a particular gene if any of these predictions were met: High/Moderate impact (VEP), possibly/probably damaging (PolyPhen-2), deleterious/deleterious_low_confidence (SIFT), or if the gene had less then 2 copies. The number of missense mutations per sample was calculated and each sample was classified as having a "high" or "low" mutation burden based on the observed bimodal distribution of missense mutation count across all samples.


1. Dupuy AJ, Rogers LM, Kim J, Nannapaneni K, Starr TK, Liu P*, et al.* A modified sleeping beauty transposon system that can be used to model a wide variety of human cancers in mice. *Cancer research* 2009 Oct 15; **69**(20)**:** 8150-8156.

2. Temiz NA, Moriarity BS, Wolf NK, Riordan JD, Dupuy AJ, Largaespada DA*, et al.* RNA sequencing of Sleeping Beauty transposon-induced tumors detects transposon-RNA fusions in forward genetic cancer screens. *Genome research* 2016 Jan; **26**(1)**:** 119-129.

3. Sarver A. Identification of Cancer Genes Based on De Novo Transposon Insertion Site Analysis Using RNA and DNA Sequencing. *Methods Mol Biol* 2019; **1907:** 73-79.

4. Li H, Durbin R. Fast and accurate short read alignment with Burrows-Wheeler transform. *Bioinformatics* 2009 Jul 15; **25**(14)**:** 1754-1760.

5. E G, G M. Haplotype-based variant detection from short-read sequencing. *arXiv* 2012; **1207:** 3907.

6. Kim D, Paggi JM, Park C, Bennett C, Salzberg SL. Graph-based genome alignment and genotyping with HISAT2 and HISAT-genotype. *Nat Biotechnol* 2019 Aug; **37**(8)**:** 907-915.

7. Liao Y, Smyth GK, Shi W. featureCounts: an efficient general purpose program for assigning sequence reads to genomic features. *Bioinformatics* 2014 Apr 1; **30**(7)**:** 923-930.

8. Robinson MD, McCarthy DJ, Smyth GK. edgeR: a Bioconductor package for differential expression analysis of digital gene expression data. *Bioinformatics* 2010 Jan 1; **26**(1)**:** 139-140.

9. Subramanian A, Tamayo P, Mootha VK, Mukherjee S, Ebert BL, Gillette MA*, et al.* Gene set enrichment analysis: a knowledge-based approach for interpreting genome-wide

expression profiles. *Proceedings of the National Academy of Sciences of the United States of America* 2005 Oct 25; **102**(43)**:** 15545-15550.

### Supplemental tables and figures

**Supplemental Table 1.** The conserved gene lists from the GCESS analysis in an excel file.

**Supplemental Table 2.** Patient sample data for the samples from the reverse phase proteomics samples in an excel file.

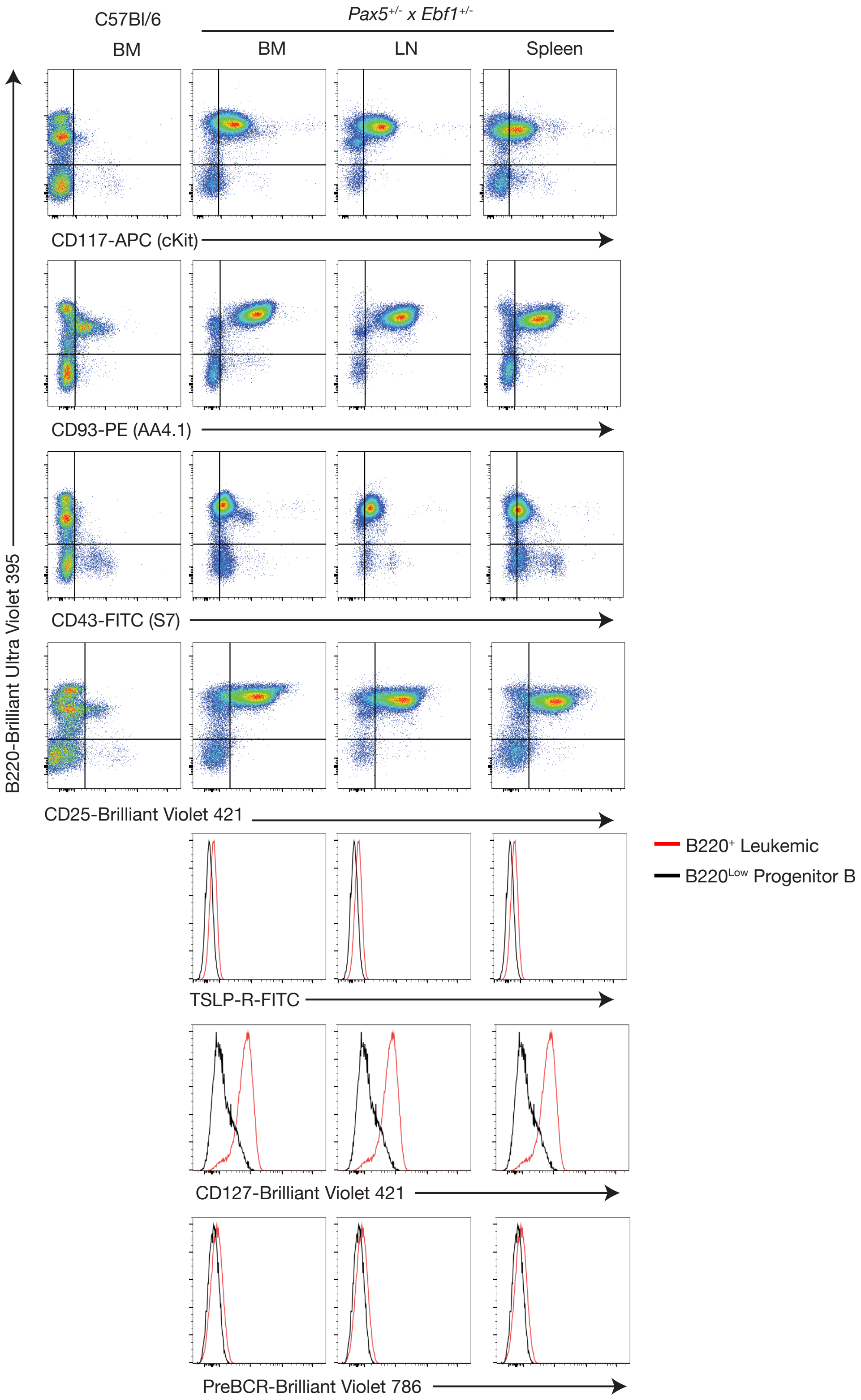


208

209 **Supplemental Figure 1.** Phenotypic characterization of progenitor B cells in *Pax5*$^{+/-}$ *x*

*Ebf1*$^{+/-}$ leukemia. Flow cytometric analysis of bone marrow cells from *Pax5*$^{+/-}$ *x Ebf1*$^{+/-}$ leukemic mice. Representative flow cytometric analysis of B220, CD117, CD93(AA4.1), CD43(S7), CD25, TSLP-R, CD127, and Pre-BCR expression on bone marrow cells from control C57Bl/6 mice or bone marrow (BM), lymph node (LN) or spleen cells from *Pax5*$^{+/-}$ *x Ebf1*$^{+/-}$ leukemic mice is shown. Doublets were gated out and a lymphocyte gate was set based on side and forward scatter properties. All gates shown are based on bone marrow isolated from control C57Bl/6 mice.

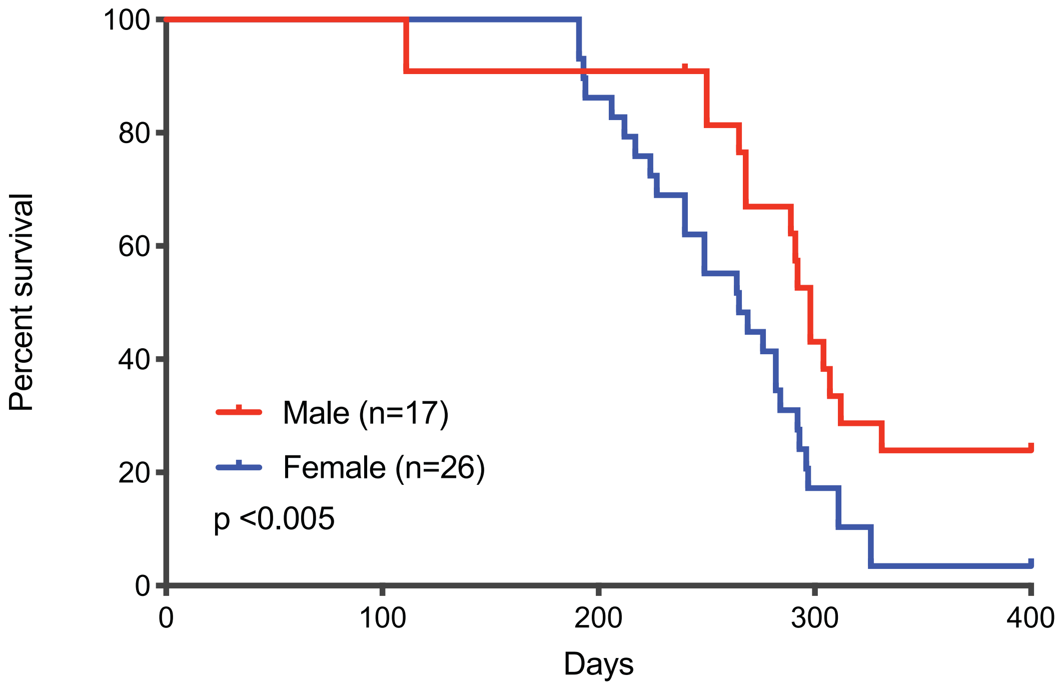


**Supplemental Figure 2.** Influence of gender on survival in *Pax5*$^{+/-}$ *x Ebf1*$^{+/-}$ leukemic mice. Kaplan-Meier survival analysis of mice of the indicated sex.

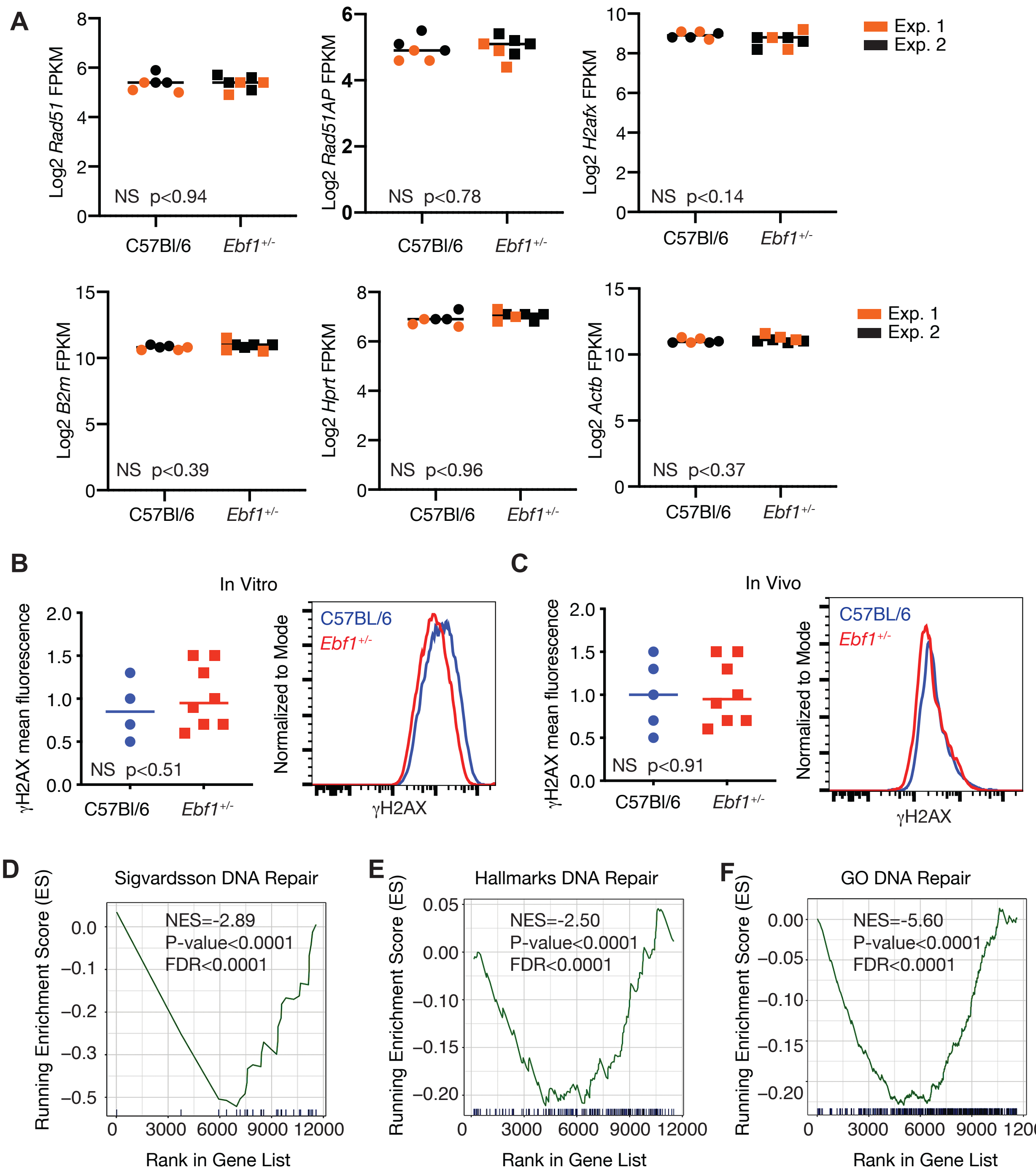


**Supplemental Figure 3.** The role of DNA repair in *Pax5*$^{+/-}$ x *Ebf1*$^{+/-}$ leukemic mice. **a** Quantitation of DNA repair genes suggested by Prasad et al to be altered in *Ebf1*$^{+/-}$ progenitor B cells and housekeeping genes. Log2 transformed FPKM values from WT (circle, n=6), *Ebf1*$^{+/-}$(square filled, n=7). We combined two independent RNA-Seq experiments and marked experiment 1 (orange) and experiment 2 (black). No significance differences were observed when using an unpaired student t-test. The line represents the median value. **b** Flow cytometry analysis of γH2AX expression in C57BL/6 or *Ebf1*$^{+/-}$ progenitor B cells grown in vitro. No significance differences were observed when using an unpaired student t-test. The graph includes samples analyzed in 3 independent experiments and the flow graph is representative of samples from all experiments. The line represents the median value. **c** Flow cytometry analysis of γH2AX expression in vivo in C57BL/6 or *Ebf1*$^{+/-}$ progenitor B cells. No significance differences were observed when using an unpaired student t-test. The graph includes samples analyzed in 3 independent experiments and the flow graph is representative of samples from all experiments. The line represents the median value. **d** Gene set enrichment (GSEA) plot comparing enrichment of genes from our ranked gene list of *Ebf1*$^{+/-}$ vs. WT data to the DNA repair gene set curated from Prasad et al., NES=-2.89, Nominal P-value < 0.0001, FDR < 0.0001 **e** GSEA plot depicting enrichment of genes from our ranked gene list compared to the Hallmark DNA repair gene set from MSigDb (M5898) NES=-2.50, Nominal P-value < 0.0001, FDR < 0.0001 **f** GSEA plot depicting enrichment of genes from our rank gene list compared to the GO DNA repair gene list from MSigDb (M12606) NES=-5.60, Nominal P-value < 0.0001, FDR < 0.0001.

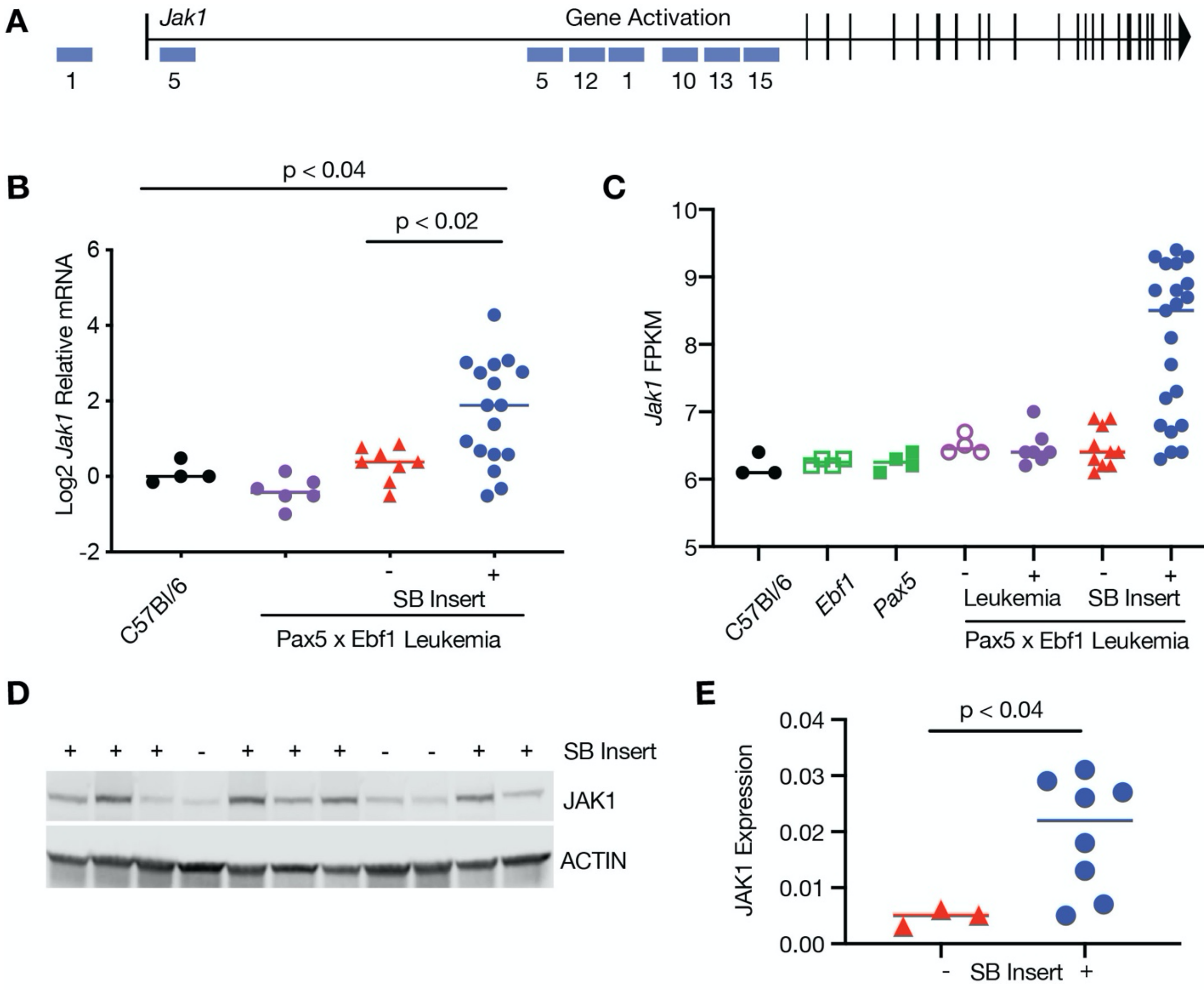


**Supplemental Figure 4.** Increased Expression of *Jak1* in leukemia. **a** Map of common insertion sites in the *Jak1* gene. **b** Quantitative Real Time PCR (qRT-PCR) for *Jak1*normalized to *Actin* in progenitor B cells isolated from the bone marrow of WT (black, n=4) mice, and leukemic cells isolated from the lymph nodes of *Pax5*$^{+/-}$ *x Ebf1*$^{+/-}$ (purple, n=6) and SB *Pax5*$^{+/-}$*x Ebf1*$^{+/-}$ mice. The samples from the SB *Pax5*$^{+/-}$*x Ebf1*$^{+/-}$ mice were split between those with (blue, n=17) or without (red, n=8) an SB insertion in the *Jak1* locus. The normalized values were log2 transformed and an ordinary one-way ANOVA with Holm-Sidak's multiple comparison test was used to determine significance. The line represents the median value. **c** Log2 transformed FPKM values from WT (black filled, n=3), *Pax5*$^{+/-}$(green filled, n=4), *Ebf1*$^{+/-}$(green open, n=4), *Pax5*$^{+/-}$ *x Ebf1*$^{+/-}$ pre-leukemic (purple open, n=4), *Pax5*$^{+/-}$ *x Ebf1*$^{+/-}$ leukemic (purple filled, n=7), and SB *Pax5*$^{+/-}$ *x Ebf1*$^{+/-}$ leukemic samples with (blue filled, n=21) or without (red filled, n=10) a transposon insertion in *Jak1* locus. A Kruskal-Wallis test with Dunn's multiple comparison test was used to test for significance. The line represents the median value. **d** Western blot analysis showing increased expression of JAK1. The + or - indicates the presence or absence of a SB transposon insertion in each representative sample. **e** Plotted ratio of JAK1 to actin from the western blot in panel d. Samples were plotted according to transposon insert status where those samples without a transposon insert are red (n=3) and those with a transposon insert are blue (n=8). Significance was determined using an unpaired student t-test. The line represents the median value.

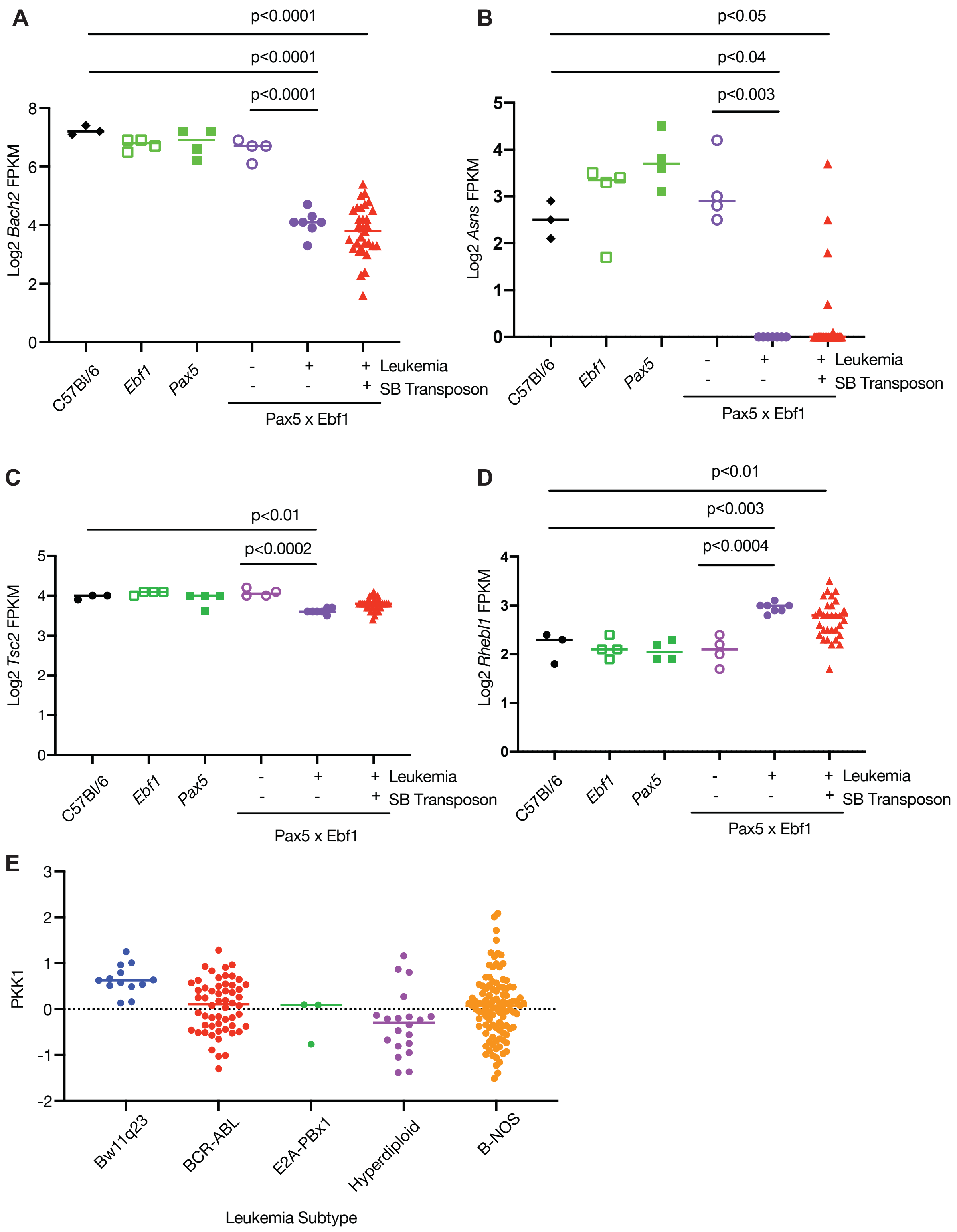
A
p<0.0001
p<0.0001
p<0.0001
Log2 Bach2 FPKM
8
6
4
2
0
C57Bl/6
Ebf1
Pax5
- + + Leukemia
- - + SB Transposon
Pax5 x Ebf1
B
p<0.05
p<0.04
p<0.003
Log2 Asns FPKM
5
4
3
2
1
0
C57Bl/6
Ebf1
Pax5
- + + Leukemia
- - + SB Transposon
Pax5 x Ebf1
C
p<0.01
p<0.0002
Log2 Tsc2 FPKM
5
4
3
2
1
0
C57Bl/6
Ebf1
Pax5
- + + Leukemia
- - + SB Transposon
Pax5 x Ebf1
D
p<0.01
p<0.003
p<0.0004
Log2 Rhebl1 FPKM
4
3
2
1
0
C57Bl/6
Ebf1
Pax5
- + + Leukemia
- - + SB Transposon
Pax5 x Ebf1
E
PKK1
3
2
1
0
-1
-2
Bw11q23
BCR-ABL
E2A-PBx1
Hyperdiploid
B-NOS
Leukemia Subtype

**Supplemental Figure 5.** Gene Expression changes in *Pax5*$^{+/-}$ *x Ebf1*$^{+/-}$ leukemia. **a** Log2 transformed FPKM values from WT (black filled, n=3), *Pax5*$^{+/-}$(green filled, n=4), *Ebf1*$^{+/-}$(green open, n=4), *Pax5*$^{+/-}$ *x Ebf1*$^{+/-}$ pre-leukemic (purple open, n=4), *Pax5*$^{+/-}$ *x Ebf1*$^{+/-}$ leukemic (purple filled, n=7), and SB *Pax5*$^{+/-}$ *x Ebf1*$^{+/-}$ leukemic (red filled, n=31) samples in the *Bach2* locus. An ordinary one-way ANOVA test with Holm-Sidak's multiple comparison test was used to test for significance. The line represents the median value.

**b** Log2 transformed FPKM values from WT (black filled, n=3), *Pax5*$^{+/-}$(green filled, n=4), *Ebf1*$^{+/-}$(green open, n=4), *Pax5*$^{+/-}$ *x Ebf1*$^{+/-}$ pre-leukemic (purple open, n=4), *Pax5*$^{+/-}$ *x Ebf1*$^{+/-}$ leukemic (purple filled, n=7), and SB *Pax5*$^{+/-}$ *x Ebf1*$^{+/-}$ leukemic (red filled, n=31) samples in the *Asns* locus. A Kruskal-Wallis test with Dunn's multiple comparison test was used to test for significance. The line represents the median value. **c** Log2 transformed FPKM values from WT (black filled, n=3), *Pax5*$^{+/-}$(green filled, n=4), *Ebf1*$^{+/-}$ (green open, n=4), *Pax5*$^{+/-}$ *x Ebf1*$^{+/-}$ pre-leukemic (purple open, n=4), *Pax5*$^{+/-}$ *x Ebf1*$^{+/-}$ leukemic (purple filled, n=7), and SB *Pax5*$^{+/-}$ *x Ebf1*$^{+/-}$ leukemic (red filled, n=31) samples in the *Tsc2* locus. A Kruskal-Wallis test with Dunn's multiple comparison test was used to test for significance. The line represents the median value. **d** Log2 transformed FPKM values from WT (black filled, n=3), *Pax5*$^{+/-}$(green filled, n=4), *Ebf1*$^{+/-}$(green open, n=4), *Pax5*$^{+/-}$ *x Ebf1*$^{+/-}$ pre-leukemic (purple open, n=4), *Pax5*$^{+/-}$ *x Ebf1*$^{+/-}$ leukemic (purple filled, n=7), and SB *Pax5*$^{+/-}$ *x Ebf1*$^{+/-}$ leukemic (red filled, n=31) samples in the *Rhebl1* locus. An ordinary one-way ANOVA test with Holm-Sidak's multiple comparison test was used to test for significance. The line represents the median value. **e** PDK1 expression of B-ALL patients split by leukemic subset Bw11q23 (blue, n=13), BCR-ABL (red, n=56), E2A-PBX1 (green, n=3), Hyperdiploid (purple, n=20) and B-NOS (orange, n=105). Total PDK1 levels were graphed for each leukemic subtype.
